# Quantum-classical hybrid approach for codon optimization and its practical applications

**DOI:** 10.1101/2024.06.08.598046

**Authors:** You Kyoung Chung, Dongkeun Lee, Junho Lee, Jaehee Kim, Daniel K Park, Joonsuk Huh

## Abstract

Codon optimization is crucial for gene expression in heterologous hosts with varying genetic codes and codon usage, potentially resulting in enhanced protein expression and stability. Traditionally, the codon optimization problem has been solved using classical numerical techniques; however, with recent advancements, quantum algorithms deployed on quantum computers have been adopted for this purpose. This study proposes a codon sequence search protocol tailored to host preferences. Specifically, codon optimization is formulated as a constrained quadratic binary problem and solved using a quantum-classical hybrid approach, integrating quantum annealing with the Lagrange multiplier method. The proposed methodology is then applied to two real-world scenarios: optimizing the codon sequence of the severe respiratory syndrome coronavirus 2 spike protein in human hosts and insulin in *Escherichia coli (E. coli)* hosts. Finally, evaluations of several biological metrics demonstrate the effectiveness of our protocol, offering insights into the codon usage patterns governing translational efficiency and adaptation to the genetic code preferences of the host organisms.

## 1 Introduction

In all genomes, most amino acids are encoded by multiple codons, with each codon comprising a sequence of three nucleotides from the messenger RNA (mRNA) that encodes a single amino acid. Typically, codon sequences are translated into polypeptides by the anticodons of transfer RNA (tRNA) within ribosomes. Of the 64 possible triplets, only 61 codons correspond to the 20 amino acids. Among these, methionine (Met) and tryptophan (Trp) are encoded by a single codon each, whereas the other 18 amino acids are encoded by two to six different codons. This phenomenon is referred to as codon degeneracy.

Despite encoding identical amino acids, these degenerate synonymous codons are unevenly distributed across the genomes of all organisms with extremely distinct frequencies [Ikemura, 1981, Ikemura, 1985]. This phenomenon, termed as codon usage bias (CUB) [Grantham et al., 1980b, Grantham et al., 1980a], has prompted researchers to explore the relationship between variations in synonymous codon preferences and gene expressions across organisms [Lloyd and Sharp, 1991, Shields and Sharp, 1987, Bennetzen and Hall, 1982]. Notably, synonymous codons exert varying influences on translation elongation speeds, highlighting the importance of their positioning within coding sequences [Letzring et al., 2010, Hiraoka et al., 2009]. Consequently, understanding codon bias patterns is crucial for elucidating the coding variability imparted by codon selection at the level of individual genes and organisms.

The frequency bias of a codon depends on the availability of tRNA molecules that deliver appropriate amino acids to the ribosome [Mitra et al., 2016]. Researchers have discovered that each organism possesses an optimal codon that is primarily utilized for protein translation. This codon bias evolves through natural selection processes and mutations and exhibits varying translation efficiencies and accuracies across organisms [Gerashchenko et al., 2021, Xu et al., 2008]. Genes with high expression levels often prefer strongly biased codons corresponding to the most abundant tRNA species [Liu, 2020, Coghlan and Wolfe, 2000]. This is because increased tRNA availability correlates with higher the translation initiation rates and improved ribosome accessibility, potentially impacting cell division and growth rates [Andersson and Kurland, 1990]. Conversely, the utilization of rare codons restricts access to tRNAs directly involved in protein translation, thus hampering gene expression levels and protein synthesis [Rosano and Ceccarelli, 2009, Kane, 1995, Clarke and Clark, 2008]. Thust, nucleotide sequences within codons influence the structure and stability of mRNA, which is crucial for gene expression levels [Lehmann and Libchaber, 2008, Victor et al., 2019].

This relationship between codons and protein expression levels indicates that optimizing codons to match the codon usage frequencies of host cells can boost translation efficiencies. Codon optimization is thus an effective strategy for enhancing the expression levels of heterologous proteins in different organisms, as exemplified by its application in the mass production of the human insulin protein in *Escherichia coli (E. coli*.) [Redwan et al., 2008, Gupta et al., 2017]. Notably, codon optimization aims to improve the codon composition of genes based on varying criteria without altering their amino acid sequences. This approach is essential to avoid the use of rare codons and to quantify CUB. Various mathematical metrics, such as the relative synonymous codon bias (RSCU) [Sharp et al., 1988], the codon adaptation index (CAI) [Sharp and Li, 1987], and the effective number of codons (ENC) [Wright, 1990, Fuglsang, 2006], have been proposed for quantitatively analyzing CUB.

In contemporary industry practices, existing commercial programs [Koblan et al., 2018,Simmonds, 2012] rely on heuristic functions. In particular, leading global companies employ various methods such as genetic algorithms (GA) [Reeves and Rowe, 2002], simulated annealing (SA) [Kirkpatrick et al., 1983], and integer linear programming [Karp, 1972] to simultaneously obtain optimal solutions for multi-objective optimization problems [Deb and Jain, 2014, Gu et al., 2021]. These techniques sample the extensive solution space by mutating synonymous codons and propagating favorable substitutions across new sequence generations. However, these programs often adopt a sliding-window approach, which encounters limitations when analyzing full-length sequences in a single computation. This is because the problem size of codon optimization is proportional to the length of sequences and closely related to NP-hardness or NP-completeness [Lucas, 2014].

With recent advancements in quantum information science and technology, quantum computing is rapidly progressing in terms of both hardware and algorithms. Recent studies have introduced various mathematical formulations and implementations of the codon optimization problem using approaches such as quantum annealing, quantum approximate optimization algorithm and variational quantum eigensolver [Fox et al., 2021, Zhang et al., 2024]. These developments highlight the potential applications of quantum computing; however, its practical applications remain unlikely owing to the limited number of qubits and various errors in quantum computing devices, commonly referred to as noisy intermediate-scale quantum (NISQ) devices [Preskill, 2018].

To address this, the current study introduces a protocol designed to tackle practical-size codon optimization problems through a quantum-classical hybrid approach, focusing on constrained optimization. Specifically, we formulate the codon optimization problem as a constrained quadratic-binary model, wherein an objective function and constraints are established employing codon statistics databases such as codon usage and codon pair usage tables. Furthermore, the constrained optimization problems is solved utilizing the Lagrange multiplier method, wherein the Lagrangian function is solved using a combination of quantum computing and classical optimization techniques. To facilitate the generation of a full-length amino acid sequences for target proteins, we specifically select quantum annealing as the quantum computing component. Notably, quantum annealing represents a heuristic optimization process tailored to a class of combinatorial optimization problems [Apolloni et al., 1990, Kadowaki and Nishimori, 1998]. Leveraging quantum properties, this approach demonstrates potential to outperform classical computers, presenting performance advantages for specific problems [Albash and Lidar, 2018, King et al., 2021]. For analysis, we adopt certain state-of-the-art quantum annealing devices, specifically the *Advantage* computer equipped with thousands of super-conducting qubits and the hybrid constrained quadratic model (CQM) solver, developed by D-Wave Systems [D-Wave Systems Inc., 2021, Yarkoni et al., 2022]. Based on the proposed protocol, we perform numerical simulations to solve real-world problems, such as identifying the codon sequences of the spike protein of the human severe respiratory syndrome coronavirus 2 (SARS-CoV-2) preferred by human hosts, as well as the codon sequences of insulin preferred by *E. coli*. Subsequently, we validate the quality of the optimized codon sequences by computing the values of several biological metrics and comparing them with reference standards.

## 2 Materials and Methods

### 2.1 Biological Metrics and their Formulations

To explore the preferred codon usage patterns of organisms and comprehend the underlying biological phenomena, researchers typically rely on various metrics or scoring functions are conventionally employed. These metrics include the CUB, codon pair usage bias (CPUB), and the guanine and cytosine (GC) content. In this section, we express these functions in the binary and quadratic forms, equivalent to the spin-1/2 Ising Hamiltonian. This mathematical form enables these metrics to be seamlessly embedded in quantum systems employed for quantum simulations or into quantum computing devices.

#### Amino Acids and Codons

According to the definition of a codon, all combinations of the four nucleotides, adenine ”A”, uracil ”U”, used by RNA instead of thymine ”T” in DNA, G, and C–generate 64 possible codons. Barring the three stop codons (UAG, UAA, and UGA), every other codon can encode one of the 20 different amino acids. Consequently, codons exhibit degeneracy for most amino acids, implying that one amino acid can be encoded by as many as six synonymous codons. For instance, arginine ”R” can be encoded by one of the following six codons: CGU, CGC, CGA, CGG, AGA, and AGG.

For any given amino acid sequence, we consider all possible synonymous codons appearing in the sequence and assign them to each of *N* binary variables {*q*_*i*_}_1≤*i*≤*N*_. Subsequently, we define the binary variable *q*_*i*_, to be embedded into qubits in the quantum annealer. According to this definition, the variable takes a value of one if a codon at position *i* is selected (*q*_*i*_ = 1) and zero otherwise (*q*_*i*_ = 0). Importantly, per this definition, the selection of an amino acid involves choosing one of its synonymous codons, and the index *i* identifies the codon and the amino acid, as illustrated in Fig. 1. For all variables {*q*_*i*_} _1≤*i*≤*N*_, the constraints ensuring that one amino acid corresponds to only one codon can be mathematically expressed as

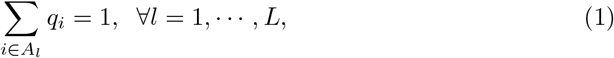

**Figure 1.**
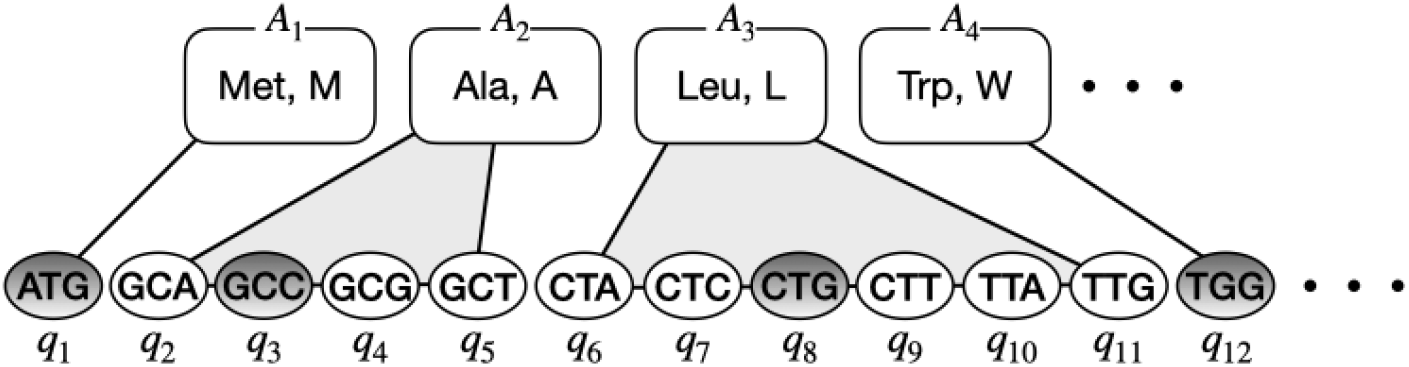
Graphical representation of all possible codons {*q*_*i*_} for the human insulin protein. Gray-colored codons depict the one amino acid-one codon selection criterion, requiring the selection of only one codon per amino acid. For instance, the second amino acid (Ala, A) has four possible synonymous codons *q*_2_, *q*_3_, *q*_4_ and *q*_5_. Among these, codon GCC is selected and mathematically expressed as *q*_3_ = 1, *q*_2_ = *q*_4_ = *q*_5_ = 0. For an amino acid sequence MALW, a vector **q** equals (1, 0, 1, 0, 0, 0, 0, 1, 0, 0, 0, 1).

where *A*_*l*_ denotes the set of synonymous codons encoding the *l*th amino acid from the available set of 20 amino acids, and *L* represents the length of the input amino acid sequence. This one amino-one codon condition is indispensable for codon optimization and will be adopted as *L* equality constraints in this study.

#### Codon Usage Bias

The CUB phenomenon is widely observed across various organisms, including bacteria, archaea, and eukaryotes [Grantham et al., 1980b, Holm, 1986]. Generally, the codon usage pattern of an organism is considered as its specific molecular signature and codon usage patterns can vary significantly across species. Consequently, comprehending CUB patterns is essential for biological studies in fields, such as genetic engineering, synthetic biology, and evolutionary studies [Ikemura, 1981, Bulmer, 1987, Sharp and Li, 1987]. The CUB phenomenon is often considered when designing synthetic genes or optimizing gene expressions to enhance the production of specific proteins in heterologous hosts [Buhr et al., 2016, Menzella, 2011]. Furthermore, evaluating CUB can offer insights into the evolutionary patterns of organisms and their adaptations to varying environmental conditions.

The empirical formula for assessing CUB relies on the concept of relative synonymous codon usage (RSCU) [Sharp and Li, 1986]. This RSCU metric compares the observed frequency of a codon with its expected frequency assuming equal codon usage. Generally, a higher RSCU value indicates a significant bias toward the usage of a specific codon, whereas a lower value suggests a less significant bias. To quantify CUB, two metrics, the CAI and ENC, are introduced based on RSCU values. Among these, the CAI quantifies the similarity between the codon usage pattern of a gene and the codon usage patterns of highly expressed genes within the same organism. Higher CAI values indicate better adaptations of codon usage patterns to the preferences of the host organism, suggesting potentially higher translation efficiencies [Sharp and Li, 1987]. Meanwhile, the ENC quantifies CUB by assesing the deviation of observed codon usage patterns from patterns involving equal usage of synonymous codons. Typically, a low ENC value indicates strong codon bias, implying that only a few codons are predominantly utilized for encoding each amino acid [Wright, 1990, Fuglsang, 2006]. Both the CAI and ENC measures offer valuable insights into translation efficiency within an organism. Typically, the CAI and ENC exhibit a negative correlation. This implies that as the CAI increases, indicating a more favorable adaptation of the codon usage pattern to the preferences of the host organism, the ENC tends to decrease, reflecting a stronger bias toward specific codons. Consequently, genes with higher CAI values are anticipated to possess lower ENC values, reflecting a more efficient and optimized utilization of codons during translation.

Within the context of this study, the CAI is employed to evaluate the qualities of optimized codon sequences. Specifically, the definition of the CAI, denoted by ℳ_CAI_, for a given codon sequence of length *L* can be expressed as

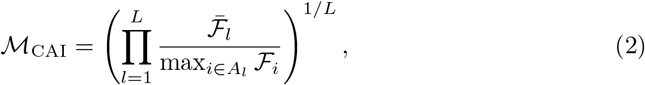

where the ℱ_*i*_ denotes the frequency of the *i*th synonymous codon of the *l*th amino acid pertaining to the *l*th codon, while the 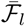 represents the frequency of appearance of the *l*th codon in a given codon sequence. The appearance or usage frequency of a codon within an organism ℱ_*i*_ can be mathematically expressed as the number of synonymous codon *i, n*_*i*_ divided by the number of the amino acid *l*, which is given by 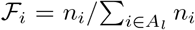 where *A*_*l*_ represents the set of the synonymous codons. For each organism, the frequency of a codon, ℱ_*i*_, can be obtained based on the average reported in the statistics of the genetic sequence database [Alexaki et al., 2019]. Notably, the CAI value ranges from zero to one: For a coding sequence comprising the most frequently used codons, the CAI value would be one.

To incorporate the concept of CUB into the framework of codon optimization, we define a scoring function that can be derived from the CAI definition. Specifically, based on the codon frequency ℱ_*i*_, the scoring function of the CUB can then be defined as a function of binary variables {*q*_*i*_},

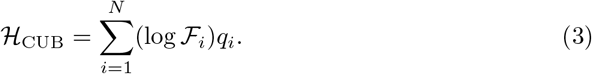

This function is derived by taking the logarithm of the CAI metric and ignoring the constant term. For the codon frequencies of an organism, solving min_*q*_ *ℋ*_CUB_ while considering the constraints specified in Eq. (1) yields the coding sequence with a CAI value of one.

#### Codon Pair Usage Bias

To examine the usage biases of individual codons, considering the usage patterns of codon pairs, particularly those appearing in close proximity within a sequence, is essential. This phenomenon involving the preferential or biased utilization of specific combinations of codons lying next to each other in an mRNA sequence is referred to as CPUB, or codon context. Notably, this bias is not random and can have functional implications. Furthermore, this bias is influenced by numerous factors including translational efficiency, mRNA stability [Yokobayashi et al., 2002, Hia et al., 2019], and the co-evolution of codon pairs. Specifically, codon pairs can affect the rate of translation elongation during protein synthesis and influence the efficiency of translation. CPUB patterns vary across diverse organisms, tissues, or environmental conditions [Parvathy et al., 2022], thus reflecting the unique evolutionary and functional characteristics of genomes. Despite being closely associated with CUB, CPUB differs in some aspects. Speficially, while CUB focuses on the usage frequencies of individual codons, CPUB considers the usage frequencies of specific pairs of codons.

Similar to the approach adopted for analyzing CUB, we can quantify CPUB by defining its scoring function. Notably, analyzing the correlation between two codons, each encoding two distinct amino acids, is essential. The usage frequency of a codon pair can be derived from the extensive set of available genetic sequences. Furthermore, individual codon information is disregarded here to solely focus on the mutual information of a codon pair. Thus, the codon pair score 𝒮_*ij*_ is deemed an appropriate quantity and is expressed as

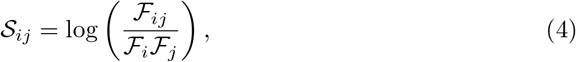

where the codon pair frequency ℱ_*ij*_ is defined as the count of the codon pair (*i, j*) divided by the count of the two amino acids encoded by the two codons (*i, j*). For a given coding sequence, the codon pair bias (CPB) is then expressed as the mean of the score for all pairs of two adjacent codons, that is, ∑_i_ 𝒮_i,i+1_*/*(*L −* 1) [Coleman et al., 2008]. Based on these mathematical expressions, the scoring function of CPUB can be defined as follows:

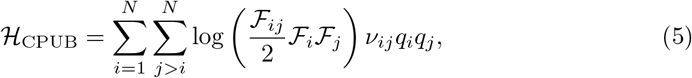

where the entries *ν*_*ij*_ of the matrix *ν* are one if *i* ∈ *A*_*l*_ and *j* ∈ *A*_*l*+1_ are satisfied and zero otherwise. Specifically, matrix *ν* ensures that the two selected codons are encoding adjacent amino acids, thus avoiding the selection of codon pairs that encode distant amino acids in the coding sequence.

#### Sequentially Repeated Nucleotide

Repeated nucleotide sequences can contribute to the structural organization of nucleic acids (DNA or RNA). A sequence having consecutively repeated nucleotides can form secondary structures, thus influencing the overall architectures of molecules. Some of these repeated nucleotide sequences can serve as functional elements within genomes, acting as regulatory elements, binding sites for proteins or other molecules, or even coding regions for functional RNA molecules [Trigiante et al., 2021]. Furthermore, repeated nucleotide sequences can influence genetic variation, potentially affecting disease susceptibility and contributing toward phenotypic diversity [Liao et al., 2023]. Thus, individual organisms display unique features and functions associated with their repeated nucleotide sequences.

In this study, we propose a scoring function for repetitive sequences, intended to serve as a penalty term in the objective function of the optimization problem [Fox et al., 2021]. Notably, this scoring function is indispensable, and its absence can adversely affect the mathematical results of codon optimization. First, we define a quantity 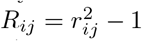 to represent the sequential repetition of nucleotides in a pair of codons (*i, j*), where *r*_*ij*_ denotes the maximal count of sequentially repeated nucleotides. Subsequently, we can then express the scoring function for sequentially repeated nucleotides as

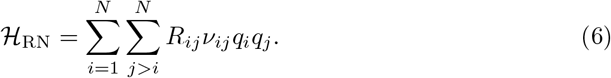

Similar to Eq. (5), matrix *ν* is introduced herein to exclusively consider the repetition of nucleotides only between two adjacent amino acids.

#### GC Concentration

In molecular biology and genetics, the GC content refers to the proportion of G and C nucleotides among the total number of nucleotides in a the DNA or RNA sequence. This metric is a crucial for the expression of various genomic features and functions [Šmarda et al., 2014]. Furthermore, variations in GC contents across organisms or genomic regions can offer insights into evolutionary processes, gene regulation, and DNA structural stability [Sharp and Lloyd, 1993,Agashe and Shankar, 2014]. Generally, high GC contents are often associated with greater DNA stability owing to the stronger bonding between G and C nucleotides compared to that between A and T nucleotides. Consequently, genes with higher GC contents in their coding regions exhibit more stable structures and exhibit different expression patterns [Rao et al., 2013]. Additionally, GC-rich regions are often linked to functional elements such as promoters and regulatory sequences [Liachko et al., 2014]. Conversely, low GC contents may indicate genome regions that are less constrained, such as non-coding regions or introns. Investigating the GC content can often insights into the structural and functional characteristics of DNA sequences [Palidwor et al., 2010, Liu et al., 2023]. Thus, we formulate the GC content using the notation of binary variables {*q*_*i*_} for codon optimization, described as

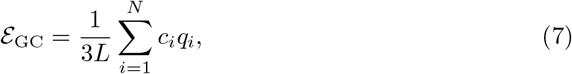

where *c*_*i*_ indicates the count of G and C bases in the *i*-th codon, and all coefficients *c*_*i*_ are determined for an input amino acid sequence.

Specifically, the GC contents exhibit notable variations, particularly at the third codon position (GC_3_ content). They tend to increase with the length of the coding sequence. Conversely, higher AT contents are observed in development-related genes [Pozzoli et al., 2008]. Therefore, in addition to the overall GC content, we also analyze the GC_3_ content as a mathematical index of codon optimization.

### 2.2 Solving Codon Optimization

Since the advent of the NISQ era, diverse quantum-classical hybrid approaches have been developed to solve combinatorial optimization problems [Preskill, 2018, Bharti et al., 2022, Cerezo et al., 2021]. Specifically, when solving constrained optimization problems, two primary methods are commonly adopted: one involving the addition of a penalty term to the objective function and another involving the Lagrange multiplier. Among these, But, the latter approach involving Lagrangian functions is significantly more beneficial for NISQ devices, as the penalty term often generate the undesired long-range interaction terms or full connectivity within NISQ devices. For instance, a linear equality constraint such as 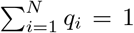 generates a penalty term involving quadratic terms such as *q*_1_*q*_*N*_. This can result in long-range interactions between distant qubits, whose high-fidelity implementation using NISQ devices is often challenging. In this section, we first formulate the codon optimization problem as a CQM, incorporating an objective function and constraints by appropriately integrating the aforementioned scoring functions. Sequentially, we provide a concise overview of both quantum annealing and the Lagrange multiplier method, each constituting a separate component of the quantum-classical hybrid approach. Finally, we outline the procedure of the quantum-classical hybrid protocol designed to address codon optimization.

#### Constrained Optimization Problems

First, we establish the mathematical forms of the objective function and constraints of codon optimization. Based on the scoring functions delineated in the previous section, we define the objective function as a linear combination of the CUB (Eq. (3)), CPUB (Eq. (5)), and sequentially repeated nucleotide (Eq. (6)) functions, as follows:

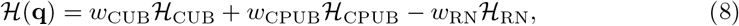

where *ω*_CUB_, *ω*_CPUB_, and *ω*_RN_ represent the non-negative weight parameters of each scoring function. These weight parameters can be adjusted depending on the relative significance of each scoring function, which will later be employed to tune the CAI and CPB values of the resulting codon sequence in the proposed protocol. Next, we set the GC content Eq. (7) as the inequality constraint to cover a specific range of the GC content. Furthermore we adopt the one amino-one codon selection (Eq. (1)) criterion as the equality constraint. Collectively, the constrained optimization problem is modeled as follows:

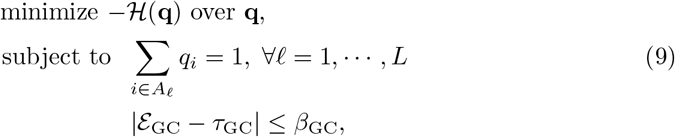

where *τ*_GC_ denotes the target value of the average GC content, and *β*_GC_ represents the tolerance of the average GC content. Notably, the objective function in Eq. (9) corresponds to the negation of ℋ(**q**), defined in Eq. (8). This adjustment is attributed to the fact that most NISQ algorithms aim to minimize the objective function rather than maximize it.

#### Quantum Annealing and Lagrange Multiplier Method

Quantum annealing is a well-established and valuable technique for solving combinatorial optimization problems [Apolloni et al., 1990, Kadowaki and Nishimori, 1998, Hauke et al., 2020, Yarkoni et al., 2022]. Unlike universal gate-based quantum computation, the range of problems that can be solved by the quantum annealing technique is limited by its underlying principles and the characteristics of quantum hardware. Nonetheless, recent studies have demonstrated that quantum annealers can solve large-scale practical problems compared to gate-based quantum computers can handle [Albash and Lidar, 2018, King et al., 2021]. Consequently, we adopted quantum annealing as it is better suited to our formulation of the codon optimization problem.

At its core, quantum annealing seeks to maintain the ground state of a Hamiltonian during time evolution. This time evolution process, extending from *t* = 0 to *t* = *T*, is governed by a time-dependent Hamiltonian, expressed as

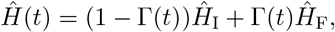

where *Γ*(*t*) is a continuous function representing the annealing schedule with *Γ*(0) = 0 and *Γ*(*T*) = 1. Furthermore, *Ĥ*_I_ represents the initial Hamiltonian, whose ground state is easy to prepare, and *Ĥ*_F_ is the final Hamiltonian, whose ground state encodes the solution to the target optimization problem. If the quantum system evolves sufficiently slowly, beginning from the ground state of *Ĥ*_I_, then the system remains in the ground state of *Ĥ* (*t*) until reaching the final Hamiltonian *Ĥ*_F_ at *t* = *T*. Typically, the initial Hamiltonian is selected as 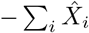 with the Pauli *X*_i_ operator acting on the *i*th qubit and the final Hamiltonian is of the Ising type, 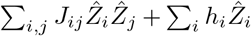 with the Pauli *Z*_*i*_ operator.

The Lagrange multiplier method is a widely adopted for solving constrained optimization problems. This method utilizes Lagrange multipliers to construct a Lagrangian function, which integrates constraints into the objective function. Based on Eq. (9), the Lagrangian function ℒ is represented as

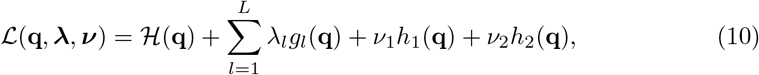

where 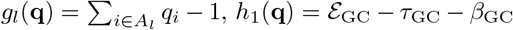 and *h*_2_(**q**) = −*ε*_GC_ + *τ*_GC_ − *β*_GC_. Here, ***λ*** = (*λ*_1_, · · ·, *λ*_*L*_) and ***ν*** = (*ν*_1_, *ν*_2_) denote Lagrange multipliers. According to the duality principle, the dual function of Eq. (9) is 𝒟(***λ, ν***) ≡ min_**q**_ **ℒ**(**q, *λ, ν***) for any ***λ*** and ***ν***. The minimum solution of Eq. (9) can be obtained by solving the dual problem, max_***λ***,***ν***_ 𝒟(***λ, ν***) subject to ***ν*** ≥ 0. Here, the dual function can be determined through quantum computing techniques dedicated to identifying the ground state of a Hamiltonian, such as quantum annealing. Meanwhile, the dual program can be handled using classical methods, allowing for a hybrid approach. Previous studies have introduced several quantum-classical hybrid algorithms embodying this philosophy [Ronagh et al., 2016, Karimi and Ronagh, 2017, Ohzeki, 2020, Djidjev, 2023, Gabbassov et al., 2023]. In this study, we specifically employ the hybrid CQM solver, developed by the D-Wave Systems, to tackle constrained optimization problems with a scale larger than those typically solved using single quantum or single classical computation alone [D-Wave Systems Inc., 2021].

#### Quantum-Classical Hybrid Codon Optimization Protocol

Below, we outline the overall procedure of the protocol adopted in this study. The primary objective of the developed protocol is to identify a coding sequence that is favored by a host organism when replicating the target amino acid sequence of another organism. The specific steps of this protocol are outlined below (See Fig. 2).

**Figure 2.**
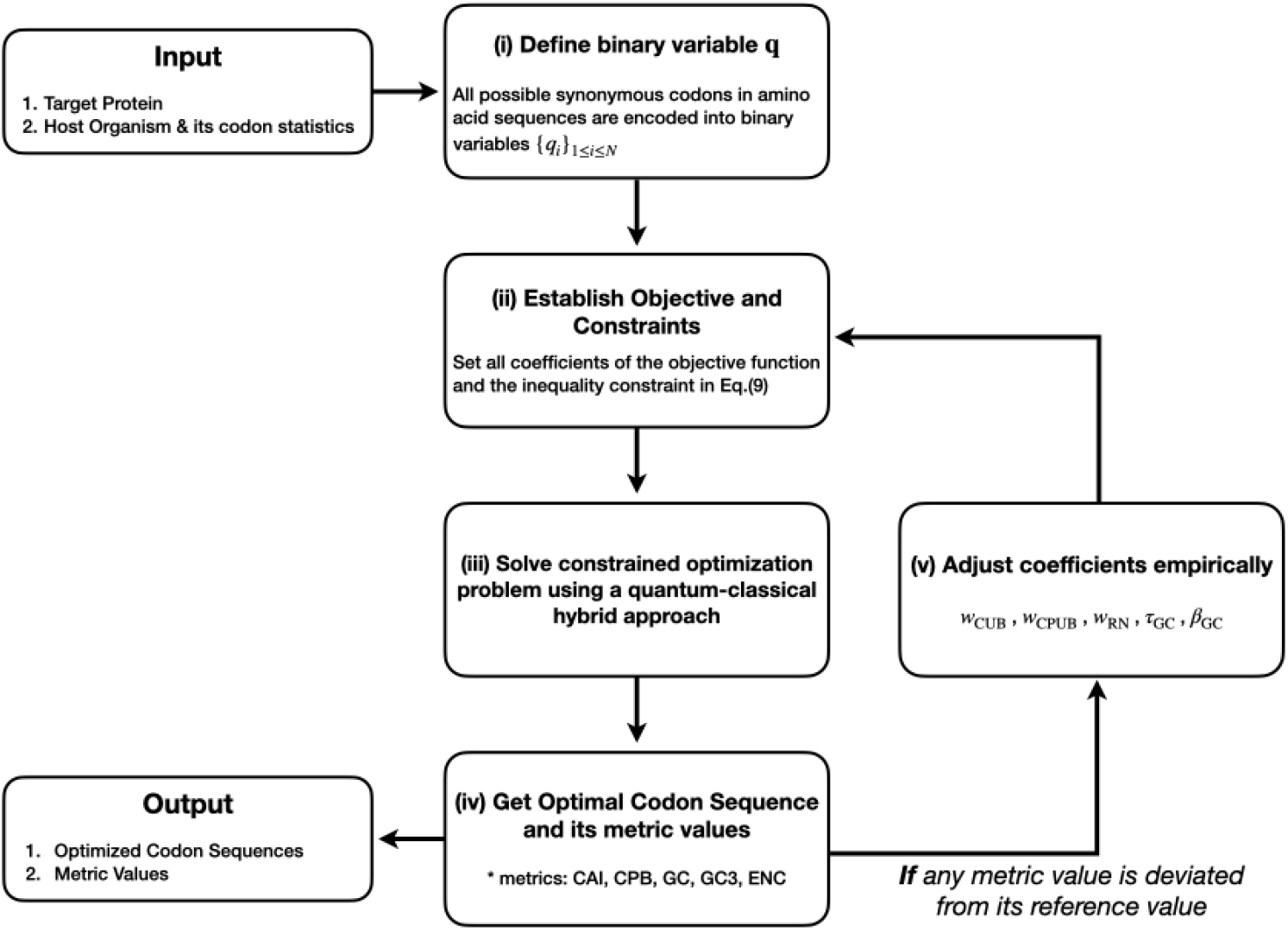
Protocol for solving the codon optimization problem using the quantum-classical hybrid approach

1. Select a target protein for expression within the host organism. Convert each amino acid *A*_*l*_ from the protein sequence into a set of binary variables (see Fig. 1) representing all possible synonymous codons of the amino acid.
2. Define the objective function and the constraints according to Eq. (9) for the given the amino acid sequence of the target protein and the host organism. Obtain the codon statistics of the host organism from available genetic data information [Alexaki et al., 2019]. These statistical data list the coefficients of several scoring functions such as the codon frequency appearing in Eq. (3) and codon pair frequency appearing in Eq. (5). Initially, set all weights *w*_CUB_, *w*_CPUB_, and *w*_RN_ in Eq. (8) to one, *τ*_GC_ to the average GC content of the host organism, and *β*_GC_ to the desired tolerance.
3. Solve the constrained codon optimization problem, expressed in Eq. (9), using the quantum-classical hybrid approach. Notably, depending on the size and the connectivity of a given problem, quantum annealing may not always guarantee that the system will reach the ground state. Thus, from the local minima obtained through the hybrid approach, select a candidate solution that is as close as possible to the optimal solution, **q**_min_.
4. Convert the optimal binary solution **q**_min_ into its corresponding nucleotide sequence according to Fig. 1. To validate the optimal coding sequence, estimate five codon metrics: the CAI, CPB, CG, GC_3_, and ENC. Subsequently, compare these values with those estimated from the extensive genetic data of the host organism.
5. If the metric values of the optimal coding sequence do not align with the reference values obtained from the genetic data, return to step 2. Subsequently, empirically adjust the coefficients of the objective function and the inequality constraint (*w*_CUB_, *w*_CPUB_, *τ*_GC_, and *β*_GC_) and repeat steps 3 and 4 until the optimal coding sequence with the desired metric values is obtained. For instance, if the CAI value of the given optimal codon sequence **q**_opt_ exceeds the corresponding value obtained from the genetic data of the host, decrease the weight *w*_CUB_ from one to 0.8 and subsequently repeat steps 3 and 4.

## 3 Results and Discussion

We implemented the proposed protocol to solve two biologically significant real-world problems. Specifically, we deployed the proposed protocol to identify and evaluate the optimal codon sequences of the SARS-CoV-2 spike protein preferred by a human host and those of insulin preferred by *E. coli*. Notably, as defined by the objective function in the constrained quadratic–binary form (Eq. (9)), the problem size depends on the type of amino acid sequence and host organism. Figure 3 depicts a graphical representation of the problem for each use case. To obtain the optimized codon sequences for these considerably long protein sequences, we selected the D-Wave hybrid CQM solver, a highly capable commercial quantum annealer, for executing step 3 of our protocol.

**Figure 3.**
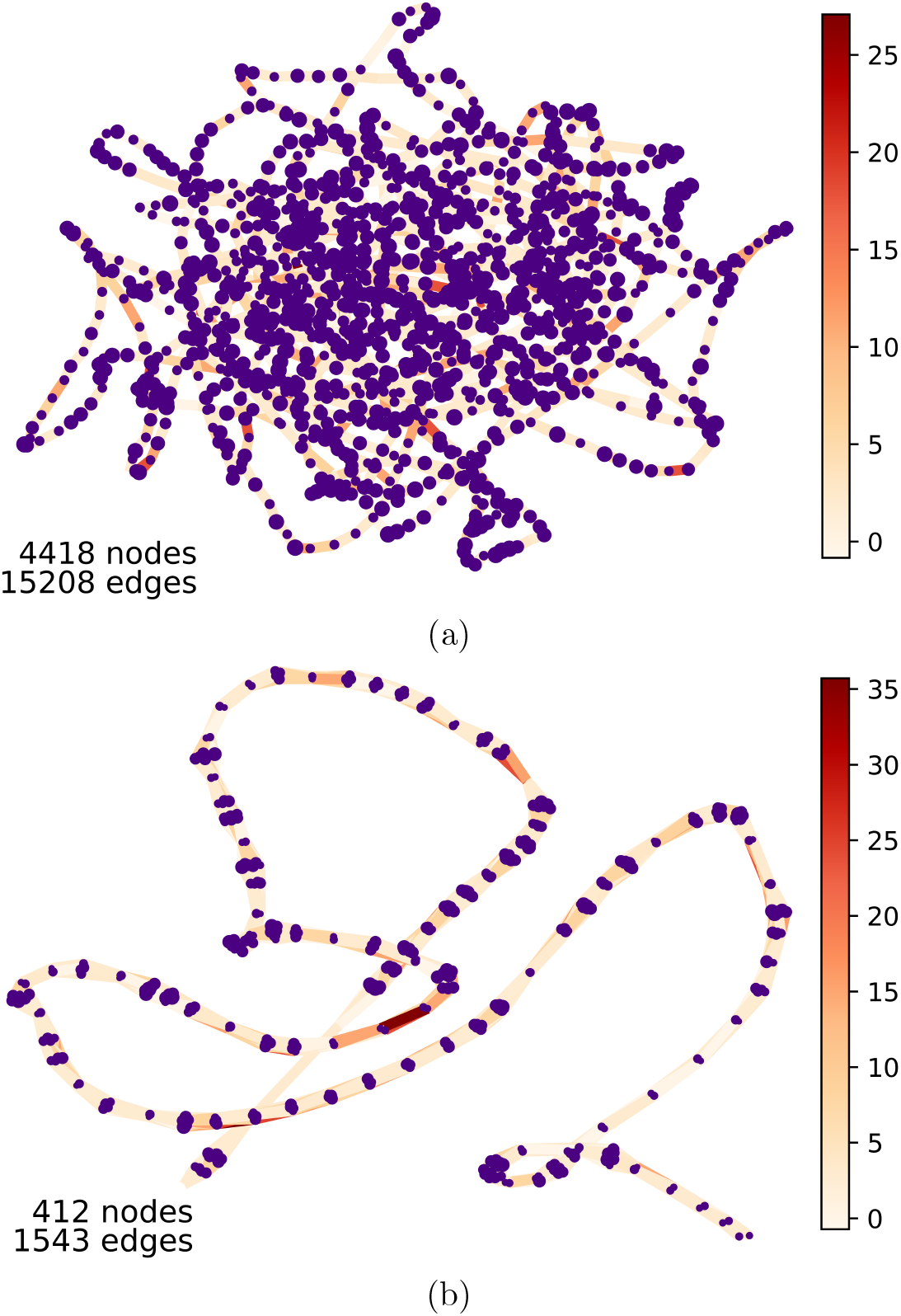
Problem graphs for two specific use cases: (a) SARS-CoV-2 spike protein in a human host and (b) insulin in *E. coli*. The vertex size and edge color indicate the coefficients of linear and quadratic terms in the objective, −ℋ(**q**) Eq. (8), respectively. These coefficients correspond to the bias and coupling strength of the Ising model.

### 3.1 SARS-CoV-2 Spike Protein

To examine the performance of our proposed protocol, we first implemented it for the codon optimization of theSARS-CoV-2 glycoprotein, employing the D-Wave hybrid CQM solver. For analysis, the complete genome sequence of SARS-CoV-2 was obtained from the National Center for Biotechnology Information [Wheeler et al., 2007]. Furthermore, the codon usage data for Homo sapiens, as the host for SARS-CoV-2, were retrieved from the codon usage database [Nakamura et al., 2000]. Notably, the target polypeptide comprised a sequence of 1,274 amino acids, which translates into a 3,822-base sequence. Upon obtaining the simulation results, the CAI, GC content, GC_3_, and ENC values of the original and optimized sequences were compared against their reference values obtained from the CAIcal 303 program [Puigbo` et al., 2007]. For CPB, we referred to the statistical distribution of CPB values for human genes reported in the literature [Coleman et al., 2008], with mean values ranging from 0.07 to 0.08. For our calculations, a mean CPB value of 0.07 was adopted as the reference for simulating the objective function. As described previously, the values of GC and GC_3_ contents vary across organisms and are tissue-specific within organisms [Li et al., 2020]. In humans, GC contents range from 20% to as high as 95%. Therefore, the reference GC contents for both the original and optimized sequences were also obtained from the literature [Li et al., 2020, Kandeel et al., 2020]. These reference values for the CAI, CPB, GC, and GC_3_ contents were utilized for the initial setup of the protocol. Meanwhile, the values of the ENC were determined based on Wright’s formula [Wright, 1990] at the end of the simulation. Following codon optimization, the CAI, GC content, GC_3_, and ENC were calculated and are summarized in Table 1.

**Table 1.** Values of the biological metrics for both the original nucleotide sequence of the SARS-CoV-2 spike glycoprotein and the corresponding optimized nucleotide sequence adapted by humans, obtained using the hybrid approach. Reference values are generated using the CAIcal software for comparison.

| Metric | Reference Values |  | Our Protocol |
| --- | --- | --- | --- |
|  | Virus host | Human host | Human host |
| CAI | 0.756 | 0.806 | 0.798 |
| CPB | 0.07 ~ 0.08 | 0.07 ~ 0.08 | 0.08 |
| GC% | 37.3 | 47.3 | 47.3 |
| GC<sub>3</sub>% | 26.7 | 53.3 | 53.2 |
| ENC | 44.2 | 36.9 | 37 |

Table 1 outlines the characteristic features of coronaviruses, including low GC content and high CAI values [Khattak et al., 2021]. The obtained results closely align with the reference values [Li et al., 2020, Kandeel et al., 2020] for all analyzed metrics, presenting only a one-digit error percentage. These findings reveal variations in CUB and base compositions between the virus and human host, despite employing the same viral sequence. Notably, the optimized sequence for the human host demonstrates a preference for GC-rich sequences [Kudla et al., 2006]. Given the high CAI value of SARS-CoV-2, the rate of gene expression is also anticipated to be high. Furthermore, higher CAI values of up to 0.8 for the human host indicate high levels of adaptation to the codon preference of the reference organism. This underscores the necessity for this pathogen to adapt to various host and environmental conditions [Šmarda et al., 2014].

#### CUB Analysis

Next, we analyzed CUB patterns in the SARS-CoV-2 spike glycoprotein genome sequence, focusing on RSCU, the nucleotide composition, and the ENC. To comprehend CUB patterns, comparisons were made with the sequences optimized for the human host. Notably, Met and Trp, each encoded by a single codon, were excluded from this analysis. Furthermore, because termination codons appear only once in each gene, they were also disregarded.

#### Relative Synonymous Codon Usage Analysis

RSCU quantifies the frequency of occurrence of different codons in the same amino acid within a given set of genes or a genome [Sharp and Li, 1987]. RSCU and the CAI are classic parameters assessing the extent of codon bias [Sharp and Li, 1986]. Typically, RSCU values greater than one indicate that a codon is used more frequently than expected, whereas RSCU values below one indicate less frequent usage than expected. Specifically, RSCU aids in assessing CUB patterns and can offer insights into gene expression, translation efficiency, and evolutionary processes [S. K. Gupta and Ghosh, 2004]. Thus, we computed the RSCU values of both the original and optimized sequences of SARS-CoV-2, and Figure 4 presents the obtained results.

**Figure 4.**
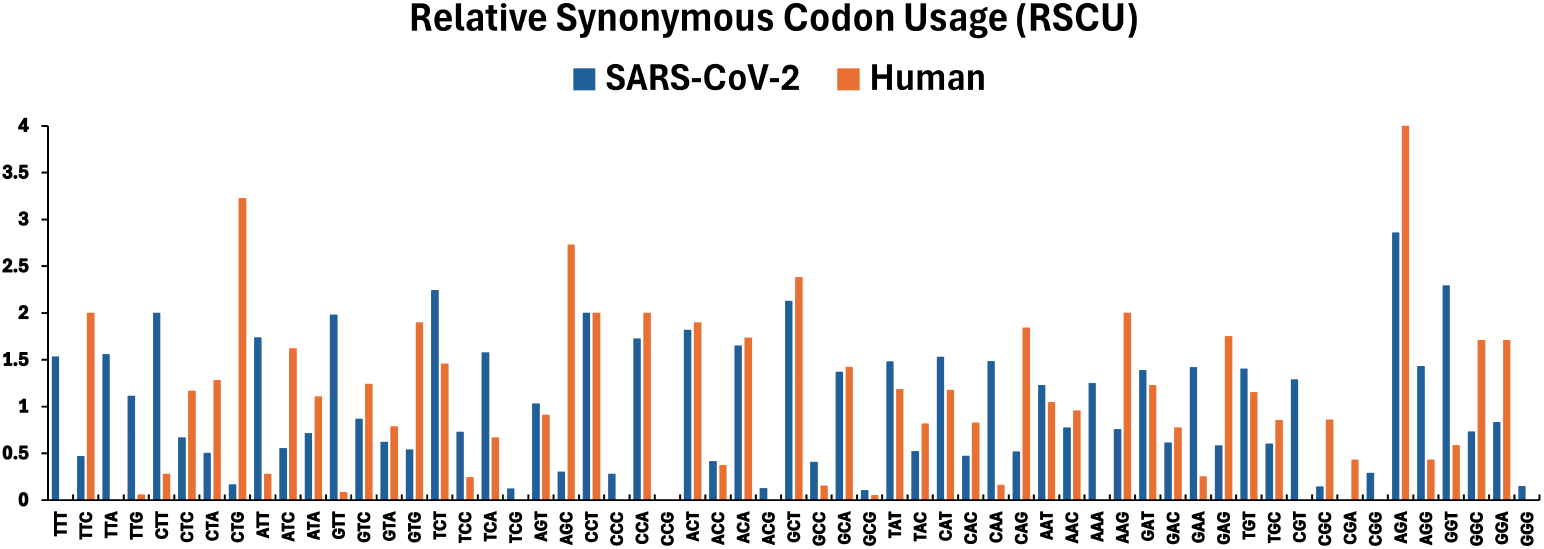
**Comparison between the RSCU values of the original SARS-CoV-2 spike glycoprotein nucleotide sequence and the corresponding optimized nucleotide sequence adapted in humans. The dark blue line represents the codons favored by SARS-CoV-2, while the orange line indicates those favored by humans**.

Figure 4 presents codons that are used more frequently or less frequently than expected. Specifically, RSCU values above 1.5 indicate codons preferred for adaptation. Our findings reveal differences between the synonymous codons preferred by the virus and those favored by the human host for the same amino acids. For instance, the virus tends to use the CTT codon for encoding the Leucine amino acid, whereas the human host preferentially selects the CTG codon, which is a G- and C-ended codon. This preference agrees with the fact that higher GC ratios necessitate high annealing and extension temperatures during the denaturation of DNA double strands into single strands, owing to the presence of three hydrogen bonds [Susp`ene et al., 2008]. Such conditions are unfavorable for the rapidly replicating virus. Consequently, the virus prefers A- and T-ended codons over G- and C-ended ones. Evidently, the AGA codon is utilized more frequently than any other codon in both the viral and human host genomes. Notably, in the SARS-CoV-2 hosted by humans, this codon is used twice as often as in the viral genome. Additionally, certain synonymous codons are utilized more or less frequently than anticipated, which is a characteristic of CUB. Notably, our findings indicate a relatively high CUB in the SARS-CoV-2 viral genome, with a preference for nearly all A or T codons [Khattak et al., 2021].

Figure 5 presents the codons preferred in both the original viral sequence and the optimized sequence. Among these, codons preferred by the SARS-CoV-2 viral gene are depicted in the outer block of the sunburst diagram. Notably, the varying selection of codons for encoding the same amino acids in different organisms suggests a propensity for overexpression or preference. Codons occupying large areas are highly preferred and represent optimal codons. Conversely, when the observed and predicted frequencies of a given codon are identical, they are depicted in the same portion of the area in the graphical representation. Conversely, an RSCU value of zero indicates the non-utilization of the given synonymous codon.

**Figure 5.**
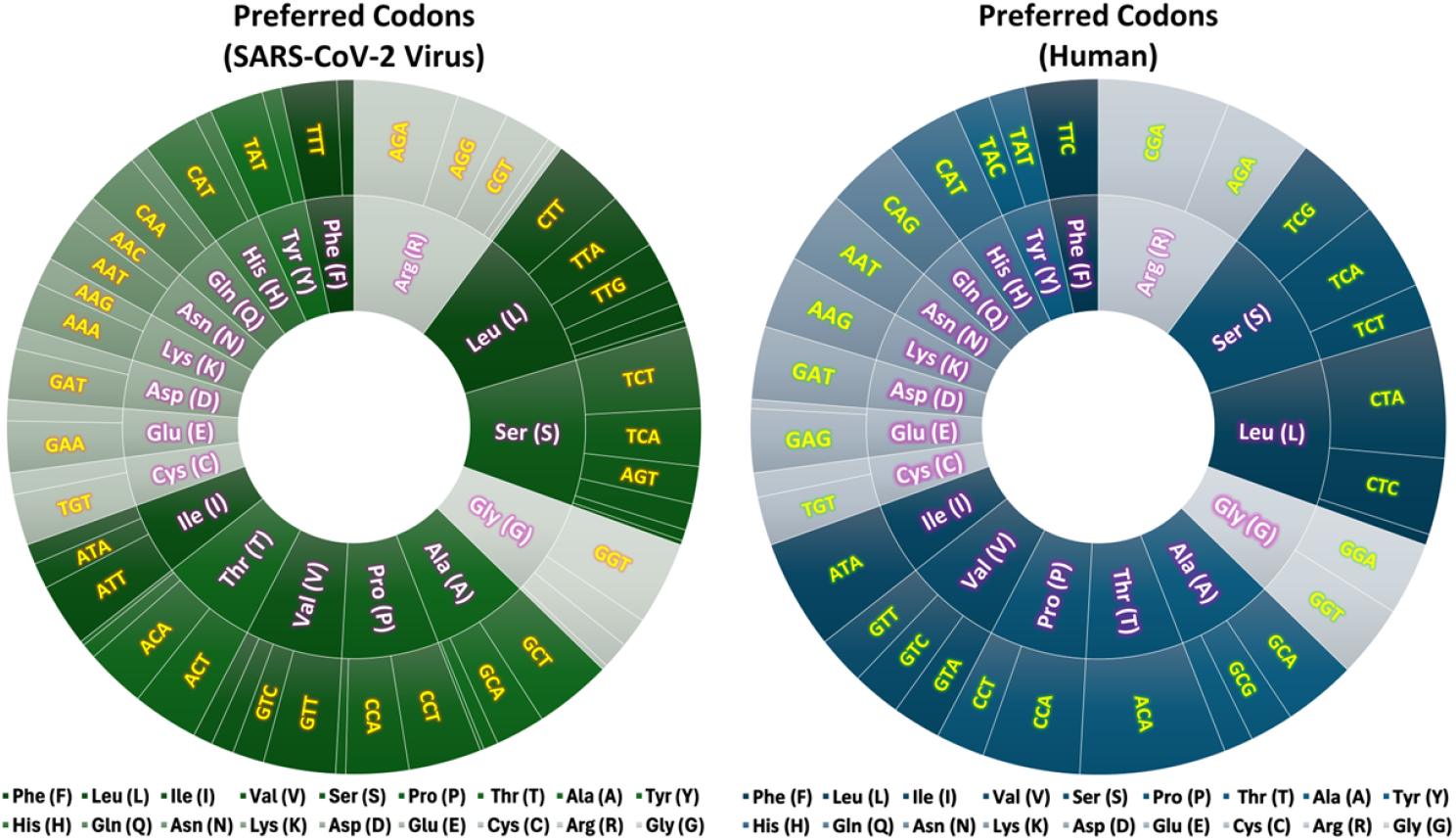
**Comparison between the codons preferred by the viral SARS-CoV-2 spike glycoprotein sequence and the optimized sequence adapted in human hosts**.

Our findings reveal differences in the selection of codons and the types of amino acids preferred by the SARS-CoV-2 virus and by the human host. For the gene encoding the SARS-CoV-2 spike protein, highly over-biased codons include CTT (Leu), ATT (Ile), GTT (Val), TCT (Ser), CCT (Pro), CCA (Pro), ACT (Thr), GCT (Ala), AGA (Arg), and GGT (Gly). Conversely, codons preferred by the human host include TTC (Phe), CTG (Leu), ATC (Ile), GTG (Val), AGC (Ser), CCT (Pro), CCA (Pro), ACT (Thr), ACA (Thr), GCT (Ala), CAG (Gln), AAG (Lys), GGC (Gly), and GGA (Gly). In the human host, the genome uses codons with higher RSCU values and GC contents more frequently. Notably, genes with higher CAI values tend to utilize more optimal codons. Given that the CAI of a gene is calculated as the geometric mean of RSCU [Sharp and Lloyd, 1993] and that RSCU is largely based on the set of genes of interest, this allows us to compare the codon biases of viral and human genes based on different aspects.

#### Nucleotide Composition Analysis

Next, we analyzed codon usage patterns by comparing the nucleotide compositions and GC contents of the SARS-CoV-2 viral genome with those of the genome adapted by the human host. Specifically, we examined the overall nucleotide compositions (A%, U%, G%, and C%) and GC contents (GC%) of both SARS-CoV-2 coding regions using the CAIcal program. Moreover, the nucleotide composition at the third codon position (A_3_, U_3_, C_3_, and G_3_) and GC_3s_ of synonymous codons were determined, and Figure 6 presents the obtained results.

**Figure 6.**
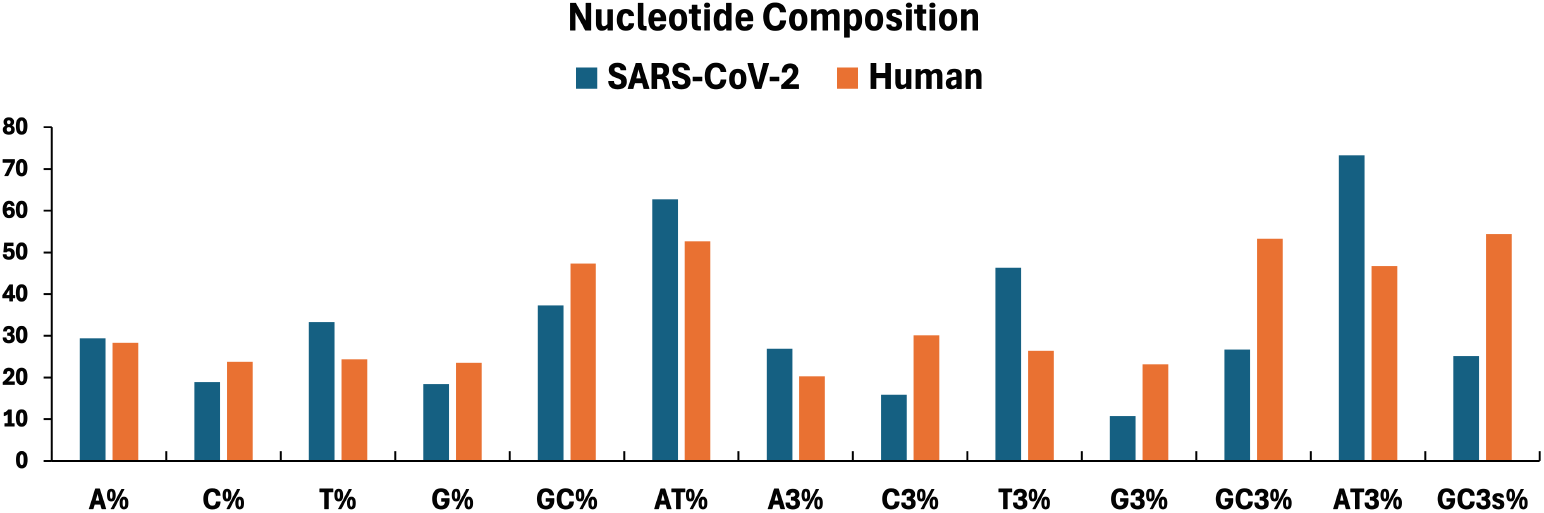
**Comparison between the nucleotide compositions of the SARS-CoV-2 spike glycoprotein sequence preferred by the virus and the codon-optimized sequence adapted by human hosts. The dark blue line denotes the nucleotides favored by SARS-CoV-2, while the orange line represents those favored by the human hosts**.

Our findings reveal that in the SARS-CoV-2 spike protein gene, T nucleotides are the most predominant (33.3%) followed by A (29.4%). SARS-CoV-2 demonstrates the lowest GC contents (37%) and the highest AT contents (63%). The nucleotide percentages in the viral genome sequence of SARS-CoV-2 are as follows: T%–A%–C%–G%. Thus, SARS-CoV-2 prefers pyrimidine-rich codons over purines. At the third codon position, T_3_ is the most frequent nucleotide, while G_3_ is the least frequent nucleotide, as presented in Fig. 6. Meanwhile, in the optimized sequence, this base composition changes to a higher GC content (53%) and lower AT content (27%). The codon composition of SARS-CoV-2 is anticipated to adapt to that of human hosts. Therefore, studying the CUB patterns of both the viral and human genomes is rational from the perspective of evolution and adaptation. Following codon optimization in the human host, the contents of G and C nucleotides significantly exceed those of the A and T nucleotides. Conversely, the nucleotide sequence of the SARS-CoV-2 viral genome presents significantly lower GC and GC_3_ contents compared to the same sequence optimized within the human host.

#### Effective Number of Codons Analysis

Further, we analyze codon usage by plotting the ENC of the SARS-CoV-2 gene against GC_3_ values. The corresponding plot is known as the NC plot [Wright, 1990]. Figure 7 displays the NC plots for the original viral RNA genome sequence of SARS-CoV-2 and the optimized codon sequence found in the human host. Similar to the previous analysis, the codons AUG (Met) and UGG (Trp) are excluded from this analysis as well.

**Figure 7.**
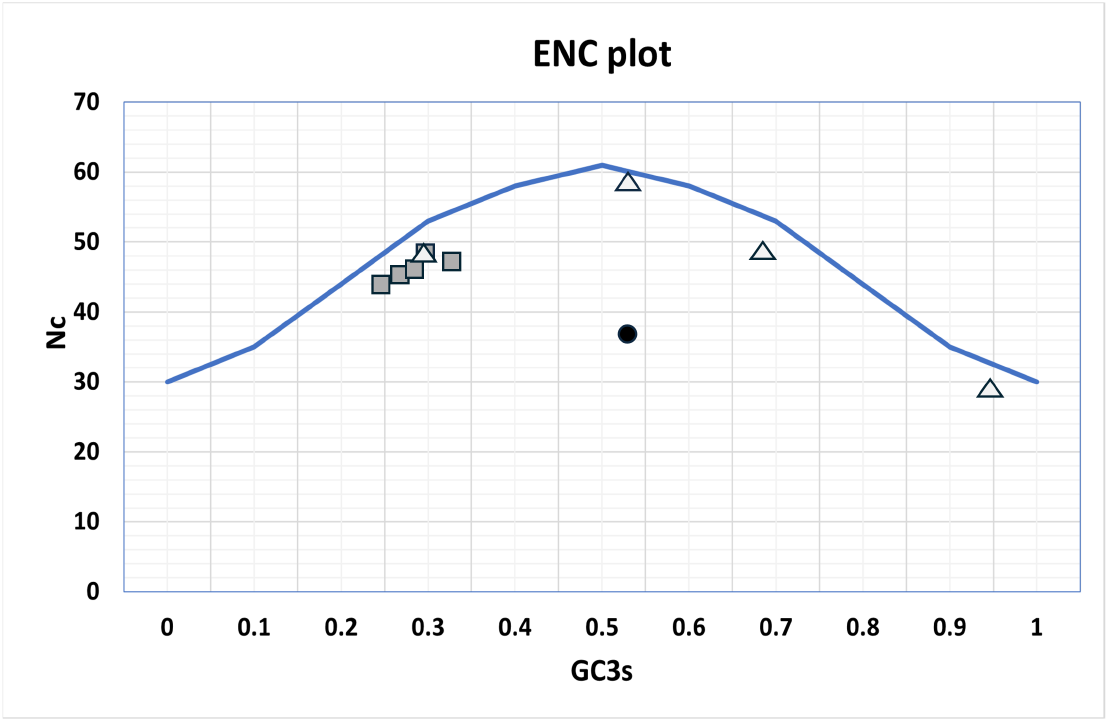
**Plot of ENC values as a function of GC**_**3**_ **for the viral SARS-CoV-2 spike glycoprotein and its codon-optimized counterpart found in human hosts. Light gray triangles represent human genes, while dark gray squares denote coronavirus genomes. The black circle indicates the simulated ENC value for the codon-optimized sequence found in the human host**.

In the figure, the continuous line represents theoretical ENC values corresponding to GC_3_ values. The data depicted by the continuous line represent NC values for some human genes (HUMAMYAS, HUMHPA2B, HUMAPOC1, and HUMHBA1) sourced from the literature [Wright, 1990]. These values are indicated by the light gray triangles. Conversely, dark gray squares correspond to the ENC and GC_3_ values of coronavirus genomes sourced from the literature [Kandeel et al., 2020, Berkhout and van Hemert, 2015]. Notably, ENC values for the viral genome range from 42 to 49, corresponding to GC_3_ contents between 0.25 and 0.35. In the figure, the ENC value corresponding to the simulated result of the codon-optimized sequence is designated with a black circle for comparison. Our findings reveal that lower ENC values indicated by the continuous line signify the translational selection of a preferred set of codons by SARS-COV-2 within the human host, similar to observations in highly expressed genes [Berkhout and van Hemert, 2015, Bennetzen and Hall, 1982, Sharp and Li, 1987]. When the ENC value lies close to or on the line, mutational pressure acts as the driving force. Conversely, when the value lies below the line, selection and mutational pressures both serve as the driving forces [Khattak et al., 2021].

As depicted, ENC values range from 20 to 61 and offer insights into CUB patterns. Specifically, a higher ENC value indicates low CUB, whereas a lower ENC value signifies strong CUB owing to the reduced number of codons utilized in protein translation [Wright, 1990]. The optimized form of the SARS-CoV-2 structural spike protein within the human host exhibits a lower ENC value of 37 than its natural counterpart within the viral genome. This suggests higher codon bias and greater gene expression efficiency of the SARS-CoV-2 structural protein within the human host. Notably, SARS-CoV-2 encodes the highest number of over-biased codons [Kandeel et al., 2020]. The plot of ENC as a function of GC_3_, depicted in Fig. 7, suggests that natural selection predominantly influences codon usage, as evidenced by the notable deviation of the ENC value from the continuous line [Berkhout and van Hemert, 2015]. The structural protein of SARS-CoV-2 exhibits a slightly lower ENC value (44) compared to other coronaviruses, including SARS CoV (45), bat SARS CoV (48), and MERS CoVs (48) [Kandeel et al., 2020]. This implies higher codon bias and enhanced gene expression efficiency of the SARS-CoV-2 structural protein. Collectively, our findings elucidate the differences in overall codon and GC usage patterns in the viral SARS-CoV-2 spike glycoprotein and its codon-optimized counterpart found in human hosts, thereby explaining the virus’s adaptation to the environment of the human host [Li et al., 2020, Šmarda et al., 2014].

### 3.2 Recombinant Protein: Insulin

For evaluations of the second use case, we assess the efficacy of the codon-optimized sequence of the human insulin gene hosted by *E. coli* in enhancing protein production. For this assessment, we compare the obtained values of the metrics with their reference values generated using the CAIcal software or sourced from the literature. Similar to the first example based on SARS-CoV-2, the proposed hybrid approach is applied to analyze the influence of codon optimization on protein expression.

#### Structure and Function of Insulin

Insulin, a crucial hormone comprising 51 amino acids, plays a vital role in regulating blood sugar levels, thus preventing conditions such as diabetes and other health issues. Dr. Frederick Banting pioneered the isolation and purification of insulin in Toronto between 1921 and 1922 [Lewis and Brubaker, 2021]. Notably, insulin can be produced on a large scale within *E. coli* bacteria owing to their robust ability to thrive on various organic substances, including sugars and proteins. Furthermore, *E. coli* bacteria are generally favored as hosts for producing recombinant proteins owing to their rapid growth, cost-effective culture conditions, and well-characterized genetics [Rosano and Ceccarelli, 2014]. Despite challenges such as protein misfolding, *E. coli* bacteria remain a popular choice for this purpose. By introducing the insulin gene into *E. coli*, these bacteria can be directed to synthesize insulin proteins, leveraging their biochemical machinery [Burgess-Brown et al., 2008]. Insulin is then synthesized as a single polypeptide, known as preproinsulin, in pancreatic beta cells [Baeshen et al., 2014]. Overall, the artificial production of insulin has been facilitated by advancements in genetic recombination technology and biotechnology. Human proinsulin, an insulin precursor comprising 110 amino acids, is located on chromosome 11 and features three regions: the A-chain, B-chain, and connecting peptide (C-peptide) [Fu et al., 2013]. The codon sequence dictates the amino acid sequence of each region, ultimately determining the structure and function of proinsulin.

#### Biological Metrics

The proposed quantum–classical hybrid protocol was adopted for codon optimization to ultimately assess and predict the influence of the human proinsulin codon sequence on protein expression efficiency in *E. coli*. Through codon optimization, we aimed to improve both the yield and quality of recombinant human insulin produced within *E. coli*, ultimately making the production process of this essential hormone in diabetes treatment more efficient and cost-effective [Burgess-Brown et al., 2008]. The simulated results of the four biological metrics are displayed in Table 2.

**Table 2.** Values of the biological metrics for both the human insulin gene and its optimized counterpart found in *E. coli* bacteria, computed using our hybrid approach. Reference values are obtained using the CAIcal program.

| Metric | Reference Values |  | Our Approach |
| --- | --- | --- | --- |
|  | Human host | *E. coli* host | *E. coli* host |
| CAI | 0.861 | 0.447 | 0.477 |
| GC% | 64.8 | 50.6 | 50.6 |
| GC<sub>3</sub>% | 80 | 38.2 | 38.2 |
| ENC | 37.5 | 32.5 | 32.5 |

As presented, the obtained results closely align with the reference values for the optimized human proinsulin gene within *E. coli*. Specifically, the initially elevated CAI values for the human insulin gene, reaching up to 0.9, decline to approximately 0.5 upon optimization within the *E. coli* host. This suggests that the *E. coli* bacteria exhibit lower CUB compared to humans, as the CAI reflects the overall bias toward the usage of specific codons within a gene or genome. A notable decrease of approximately 22% in the GC content and another decrease of 50% in GC_3_ values are observed when comparing the data corresponding to humans with those corresponding to *E. coli*. This difference underscores the distinct genetic compositions and environments of the compared organisms, which influence their codon bias patterns. In humans, the higher GC content is likely associated with a stronger preference for GC-rich codons, contributing to mRNA stability, protein folding, and gene expression regulation. This bias reflects the evolutionary history and functional constraints of the human genome. Conversely, *E. coli* bacteria, which are prokaryotic organisms, exhibit distinct patterns in the GC content and codon bias compared to humans. Notably, the lower ENC value following codon optimization in *E. coli* indicates a stronger bias toward specific codons in the genetic code of *E. coli*. This bias suggests that certain codons are preferred over others for protein synthesis in *E. coli*, potentially leading to more efficient and accurate translation processes. Overall, investigations into the codon bias patterns of humans and *E. coli* offer insights into the mechanisms of gene expression regulation and protein synthesis. By analyzing the CAI, CUB, CPB, and GC contents, researchers can gain valuable insights into the evolutionary and functional characteristics of the genes and genomes of various organisms.

#### Analysis of Codon Usage Patterns

To comprehend the non-random usage patterns of synonymous codons in *E. coli* for human insulin production, we analyzed and compared RSCU, the nucleotide composition, and the GC variation of each gene sequence across the two organisms.

Figure 8(a) depicts the nucleotide compositions of both the original and the codon-optimized human insulin gene sequences. Notably, within the human insulin gene, the G and C nucleotides are predominant, corroborating its high GC content (65%) and low AT content (35%). However, in the optimized gene sequence within *E. coli*, the GC content decreases, whereas the AT content increases. In particular, the A content increases significantly. This shift suggests that while eukaryotes such as humans prefer GC-rich codons, prokaryotes such as *E. coli* favor AT-rich codons. Bacteria, being less evolved prokaryotes, tend to adopt simpler evolutionary mechanisms [Shi et al., 2022] for rapid division and growth. Consequently, *E. coli* bacteria exhibit a preference for AT-rich codons, which require lower temperatures for annealing and extension [Obradovic et al., 2013]. Conversely, evolved eukaryotes [Lynch and Marinov, 2017] develop defense mechanisms against reactive oxygen species through sufficient ATP synthesis in their mitochondria [Phaniendra et al., 2015]. Consequently, the functions of eukaryotes, including humans, are often completely developed. Conversely, prokaryotic genomes exhibit various GC contents, ranging from 13% to 75%, enabling bacteria to adapt to diverse environments [Yoshikawa and Sueoka, 1963]. Figure 8(b) presents the varying degrees of changes observed at each base position in the nucleotide sequence. Regarding the GC content, the first and second base compositions remain relatively stable, whereas the third base composition presents a notable decrease during codon selection in the *E. coli* system. This signifies the importance of the base position, even within one codon. Generally, *E. coli* prefer A- and T-ending codons, demonstrating a noticeable variation in the third base position during codon optimization of the human insulin gene. Compared to the ENC value for the human insulin gene sequence, that for the optimized sequences preferred by *E. coli* slightly decreased. This suggests that fewer codons are being utilized to encode the same amino acid, reflecting more efficient usage of codons during translation. From this observation, we can infer that the gene sequence may have been subjected to selective conditions or evolutionary processes, favoring specific codons over others. This could be attributed to factors such as adaptation to the host’s translational machinery or optimization for efficient protein synthesis.

**Figure 8.**
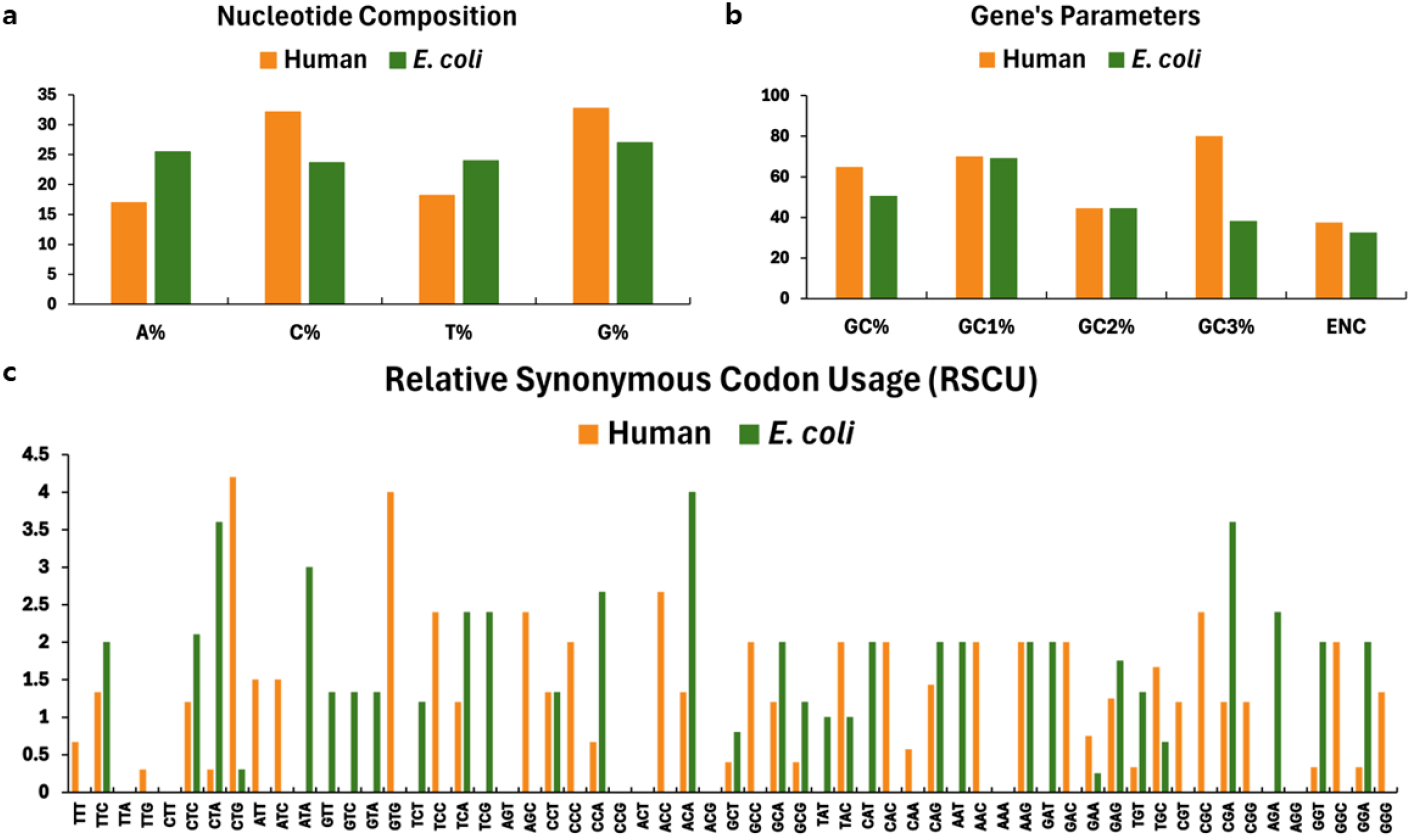
**Comparisons between the (a) nucleotide compositions; (b) numerical parameters, such as GC contents and ENC values; and (c) RSCU patterns of the human insulin gene sequence and its codon-optimized counterpart within *E. coli* bacteria. Orange lines depict the nucleotides and codons preferred by humans, while green lines represent those favored by *E. coli***.

Figure 8(c) depicts the RSCU patterns of the preferred codons within the human insulin gene. Notably, the most over-biased codons in humans (RSCU*>*2.5) include CTG (Leu) and GTG (Val), whereas in *E. coli*, these include ACA (Thr), CTA (Leu), ATA (Ile), and CGA (Arg). Figure 9 compares the preferred codons within the human insulin gene and its codon-optimized counterpart found in *E. coli*.

**Figure 9.**
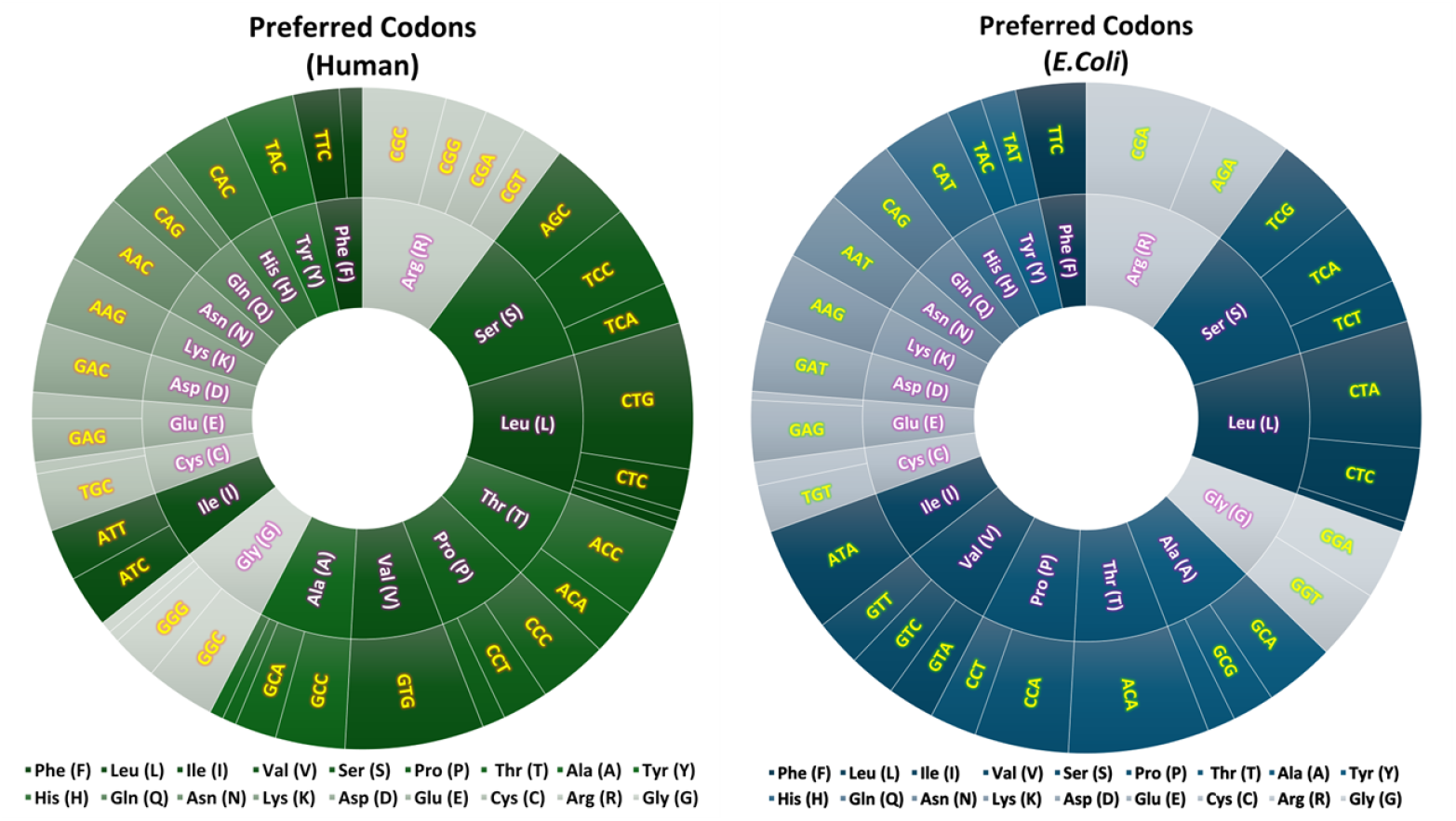
**Comparison between the preferred codons found in the human insulin gene sequence and its codon-optimized counterpart found in *E. coli* bacteria**.

We observe that while humans prefer the CTG codon, *E. coli* bacteria utilize the CTA codon more frequently for encoding the same Leucine amino acid during human insulin protein synthesis. This discrepancy highlights that a high RSCU value for a codon suggests a preference for its usage, indicating a bias in codon selection. Within *E. coli* the choices of codons encoding abundant protein species are strictly dependent on tRNA availability [Ikemura, 1981]. For less frequently utilized codons, evidence suggests co-evolution between codon usage and the concentrations of isoacceptor tRNAs. This correlation suggests that preferred codons correspond to the abundance of cognate tRNAs present within the cell [Gustafsson et al., 2004]. For instance, *E. coli* bacteria prefer TTC over TTT for encoding phenylalanine amino acid, enhancing the efficiency of the translation process improving the efficiency of the translation process [Ikemura, 1981]. Generally, genes with higher CAI values tend to employ more optimal codons. The observed differences in the CAI values of the human insulin genes hosted by humans and by *E. coli* highlight the contrasting optimal codon bias between these organisms. This discrepancy prompts a comparative analysis of the codon usage patterns evident in *E. coli* and humans, aiming to acquire insights into the evolutionary process from *E. coli* to humans. Furthermore, analyzing nucleotide distributions offers insights into the overall GC content and base pair preference within a gene, enriching our understanding of genetic diversity and evolutionary processes, thus improving future genetic engineering efforts such as insulin biosynthesis.

## 4 Conclusion

Codon bias, a widespread biological phenomenon apparent in various organisms, demonstrates significant implications and applications across diverse domains. Codon optimization is essential for maximizing gene expression efficiency. This study presents an advanced protocol based on a quantum–classical hybrid approach, integrating quantum annealing with the Lagrange multiplier method, to solve practical-size codon optimization problems formulated as constrained quadratic–binary problems. Leveraging the unique capabilities of state-of-the art quantum annealers, we apply the developed protocol to two real-world scenarios: optimizing the codon sequences of the SARS-CoV-2 spike protein preferred by human cells and the codon sequences for insulin preferred by *E. coli*. Through extensive analyses, we validate the effectiveness of our protocol by comparing the metric values of output sequences with their average values obtained from genetic databases. Our findings reveal notable differences in codon bias patterns across various species, offering valuable insights into genetic features and variability. Thus, we can conclude that codon optimization presents a powerful means for enhancing gene expression efficiency and protein production by leveraging the codon usage preferences of the host organism.

We envision that our approach will offer a promising avenue for addressing codon optimization, leveraging advancements in quantum computing technology. As highlighted in the main text, the computational cost associated with codon optimization problems depends on the type and length of the input protein sequence and the host organism. While the superiority of contemporary quantum computers over classical computers in solving various problems is currently unclear, a combination of quantum annealing and classical methods offer promising solutions, especially for tackling combinatorial optimization problems. Refining the mathematical expression of codon optimization to maintain pace with advancements in quantum computing or incorporating additional metrics into its mathematical framework may yield better quality codon sequences. This not only applies to codon optimization but also presents opportunities for transformative breakthroughs in various fields, including protein design and drug discovery.

## Acknowledgments

This work was partly supported by the Basic Science Research Program through the National Research Foundation of Korea (NRF), funded by the Ministry of Education, Science and Technology (NRF-2022M3H3A106307411, NRF-2023M3K5A1094805, and NRF-2023M3K5A109481311), Institute for Information & communications Technology Promotion (IITP) grant funded by the Korea government(MSIP) (No. 2019-0-00003, Research and Development of Core technologies for Programming, Running, Implementing and Validating of Fault-Tolerant Quantum Computing System), by the KIST Institutional Program (2E32941-24-008), and by the Yonsei University Research Fund of 2024 (2024-22-0147). This research was also partly supported by ‘Quantum Information Science R&D Ecosystem Creation’ through the National Research Foundation of Korea(NRF) funded by the Korean government (Ministry of Science and ICT(MSIT))(No. 2020M3H3A1110365) and by the National Research Foundation of Korea (NRF) grant (NRF-2022M3K2A1083890) funded by the Korea Government (MSIP).

## A Optimized Codon Sequences obtained from our protocol

### A.1 SARS-CoV-2 Spike Protein

#### 1. Protein sequence

YP 009724390.1 S [organism=Severe acute respiratory syndrome coronavirus 2] [GeneID=43740568]

MFVFLVLLPLVSSQCVNLTTRTQLPPAYTNSFTRGVYYPDKVFRSSVLHSTQDLFLPFFSNVTWFHAIHVSGTNGTKRFDNPVLPFNDGVYFASTEKSNIIRGWIFGTTLDSKTQSLLIVNNATNVVIKVCEFQFCNDPFLGVYYHKNNKSWMESEFRVYSSANNCTFEYVSQPFLMDLEGKQGNFKNLREFVFKNIDGYFKIYSKHTPINLVRDLPQGFSALEPLVDLPIGINITRFQTLLALHRSYLTPGDSSSGWTAGAAAYYVGYLQPRTFLLKYNENGTITDAVDCALDPLSETKCTLKSFTVEKGIYQTSNFRVQPTESIVRFPNITNLCPFGEVFNATRFASVYAWNRKRISNCVADYSVLYNSASFSTFKCYGVSPTKLNDLCFTNVYADSFVIRGDEVRQIAPGQTGKIADYNYKLPDDFTGCVIAWNSNNLDSKVGGNYNYLYRLFRKSNLKPFERDISTEIYQAGSTPCNGVEGFNCYFPLQSYGFQPTNGVGYQPYRVVVLSFELLHAPATVCGPKKSTNLVKNKCVNFNFNGLTGTGVLTESNKKFLPFQQFGRDIADTTDAVRDPQTLEILDITPCSFGGVSVITPGTNTSNQVAVLYQDVNCTEVPVAIHADQLTPTWRVYSTGSNVFQTRAGCLIGAEHVNNSYECDIPIGAGICASYQTQTNSPRRARSVASQSIIAYTMSLGAENSVAYSNNSIAIPTNFTISVTTEILPVSMTKTSVDCTMYICGDSTECSNLLLQYGSFCTQLNRALTGIAVEQDKNTQEVFAQVKQIYKTPPIKDFGGFNFSQILPDPSKPSKRSFIEDLLFNKVTLADAGFIKQYGDCLGDIAARDLICAQKFNGLTVLPPLLTDEMIAQYTSALLAGTITSGWTFGAGAALQIPFAMQMAYRFNGIGVTQNVLYENQKLIANQFNSAIGKIQDSLSSTASALGKLQDVVNQNAQALNTLVKQLSSNFGAISSVLNDILSRLDKVEAEVQIDRLITGRLQSLQTYVTQQLIRAAEIRASANLAATKMSECVLGQSKRVDFCGKGYHLMSFPQSAPHGVVFLHVTYVPAQEKNFTTAPAICHDGKAHFPREGVFVSNGTHWFVTQRNFYEPQIITTDNTFVSGNCDVVIGIVNNTVYDPLQPELDSFKEELDKYFKNHTSPDVDLGDISGINASVVNIQKEIDRLNEVAKNLNESLIDLQELGKYEQYIKWPWYIWLGFIAGLIAIVMVTIMLCCMTSCCSCLKGCCSCGSCCKFDEDDSEPVLKGVKLHYT

#### 2. Nucleotide sequence

NC 045512.2:21563-25384 S [organism=Severe acute respiratory syndrome coronavirus 2] [GeneID=43740568] [chromosome=]

ATGTTTGTTTTTCTTGTTTTATTGCCACTAGTCTCTAGTCAGTGTGTTAATCTTACAACCAGAACTCAATTACCCCCTGCATACACTAATTCTTTCACACGTGGTGTTTATTACCCTGACAAAGTTTTCAGATCCTCAGTTTTACATTCAACTCAGGACTTGTTCTTACCTTTCTTTTCCAATGTTACTTGGTTCCATGCTATACATGTCTCTGGGACCAATGGTACTAAGAGGTTTGATAACCCTGTCCTACCATTTAATGATGGTGTTTATTTTGCTTCCACTGAGAAGTCTAACATAATAAGAGGCTGGATTTTTGGTACTACTTTAGATTCGAAGACCCAGTCCCTACTTATTGTTAATAACGCTACTAATGTTGTTATTAAAGTCTGTGAATTTCAATTTTGTAATGATCCATTTTTGGGTGTTTATTACCACAAAAACAACAAAAGTTGGATGGAAAGTGAGTTCAGAGTTTATTCTAGTGCGAATAATTGCACTTTTGAATATGTCTCTCAGCCTTTTCTTATGGACCTTGAAGGAAAACAGGGTAATTTCAAAAATCTTAGGGAATTTGTGTTTAAGAATATTGATGGTTATTTTAAAATATATTCTAAGCACACGCCTATTAATTTAGTGCGTGATCTCCCTCAGGGTTTTTCGGCTTTAGAACCATTGGTAGATTTGCCAATAGGTATTAACATCACTAGGTTTCAAACTTTACTTGCTTTACATAGAAGTTATTTGACTCCTGGTGATTCTTCTTCAGGTTGGACAGCTGGTGCTGCAGCTTATTATGTGGGTTATCTTCAACCTAGGACTTTTCTATTAAAATATAATGAAAATGGAACCATTACAGATGCTGTAGACTGTGCACTTGACCCTCTCTCAGAAACAAAGTGTACGTTGAAATCCTTCACTGTAGAAAAAGGAATCTATCAAACTTCTAACTTTAGAGTCCAACCAACAGAATCTATTGTTAGATTTCCTAATATTACAAACTTGTGCCCTTTTGGTGAAGTTTTTAACGCCACCAGATTTGCATCTGTTTATGCTTGGAACAGGAAGAGAATCAGCAACTGTGTTGCTGATTATTCTGTCCTATATAATTCCGCATCATTTTCCACTTTTAAGTGTTATGGAGTGTCTCCTACTAAATTAAATGATCTCTGCTTTACTAATGTCTATGCAGATTCATTTGTAATTAGAGGTGATGAAGTCAGACAAATCGCTCCAGGGCAAACTGGAAAGATTGCTGATTATAATTATAAATTACCAGATGATTTTACAGGCTGCGTTATAGCTTGGAATTCTAACAATCTTGATTCTAAGGTTGGTGGTAATTATAATTACCTGTATAGATTGTTTAGGAAGTCTAATCTCAAACCTTTTGAGAGAGATATTTCAACTGAAATCTATCAGGCCGGTAGCACACCTTGTAATGGTGTTGAAGGTTTTAATTGTTACTTTCCTTTACAATCATATGGTTTCCAACCCACTAATGGTGTTGGTTACCAACCATACAGAGTAGTAGTACTTTCTTTTGAACTTCTACATGCACCAGCAACTGTTTGTGGACCTAAAAAGTCTACTAATTTGGTTAAAAACAAATGTGTCAATTTCAACTTCAATGGTTTAACAGGCACAGGTGTTCTTACTGAGTCTAACAAAAAGTTTCTGCCTTTCCAACAATTTGGCAGAGACATTGCTGACACTACTGATGCTGTCCGTGATCCACAGACACTTGAGATTCTTGACATTACACCATGTTCTTTTGGTGGTGTCAGTGTTATAACACCAGGAACAAATACTTCTAACCAGGTTGCTGTTCTTTATCAGGATGTTAACTGCACAGAAGTCCCTGTTGCTATTCATGCAGATCAACTTACTCCTACTTGGCGTGTTTATTCTACAGGTTCTAATGTTTTTCAAACACGTGCAGGCTGTTTAATAGGGGCTGAACATGTCAACAACTCATATGAGTGTGACATACCCATTGGTGCAGGTATATGCGCTAGTTATCAGACTCAGACTAATTCTCCTCGGCGGGCACGTAGTGTAGCTAGTCAATCCATCATTGCCTACACTATGTCACTTGGTGCAGAAAATTCAGTTGCTTACTCTAATAACTCTATTGCCATACCCACAAATTTTACTATTAGTGTTACCACAGAAATTCTACCAGTGTCTATGACCAAGACATCAGTAGATTGTACAATGTACATTTGTGGTGATTCAACTGAATGCAGCAATCTTTTGTTGCAATATGGCAGTTTTTGTACACAATTAAACCGTGCTTTAACTGGAATAGCTGTTGAACAAGACAAAAACACCCAAGAAGTTTTTGCACAAGTCAAACAAATTTACAAAACACCACCAATTAAAGATTTTGGTGGTTTTAATTTTTCACAAATATTACCAGATCCATCAAAACCAAGCAAGAGGTCATTTATTGAAGATCTACTTTTCAACAAAGTGACACTTGCAGATGCTGGCTTCATCAAACAATATGGTGATTGCCTTGGTGATATTGCTGCTAGAGACCTCATTTGTGCACAAAAGTTTAACGGCCTTACTGTTTTGCCACCTTTGCTCACAGATGAAATGATTGCTCAATACACTTCTGCACTGTTAGCGGGTACAATCACTTCTGGTTGGACCTTTGGTGCAGGTGCTGCATTACAAATACCATTTGCTATGCAAATGGCTTATAGGTTTAATGGTATTGGAGTTACACAGAATGTTCTCTATGAGAACCAAAAATTGATTGCCAACCAATTTAATAGTGCTATTGGCAAAATTCAAGACTCACTTTCTTCCACAGCAAGTGCACTTGGAAAACTTCAAGATGTGGTCAACCAAAATGCACAAGCTTTAAACACGCTTGTTAAACAACTTAGCTCCAATTTTGGTGCAATTTCAAGTGTTTTAAATGATATCCTTTCACGTCTTGACAAAGTTGAGGCTGAAGTGCAAATTGATAGGTTGATCACAGGCAGACTTCAAAGTTTGCAGACATATGTGACTCAACAATTAATTAGAGCTGCAGAAATCAGAGCTTCTGCTAATCTTGCTGCTACTAAAATGTCAGAGTGTGTACTTGGACAATCAAAAAGAGTTGATTTTTGTGGAAAGGGCTATCATCTTATGTCCTTCCCTCAGTCAGCACCTCATGGTGTAGTCTTCTTGCATGTGACTTATGTCCCTGCACAAGAAAAGAACTTCACAACTGCTCCTGCCATTTGTCATGATGGAAAAGCACACTTTCCTCGTGAAGGTGTCTTTGTTTCAAATGGCACACACTGGTTTGTAACACAAAGGAATTTTTATGAACCACAAATCATTACTACAGACAACACATTTGTGTCTGGTAACTGTGATGTTGTAATAGGAATTGTCAACAACACAGTTTATGATCCTTTGCAACCTGAATTAGACTCATTCAAGGAGGAGTTAGATAAATATTTTAAGAATCATACATCACCAGATGTTGATTTAGGTGACATCTCTGGCATTAATGCTTCAGTTGTAAACATTCAAAAAGAAATTGACCGCCTCAATGAGGTTGCCAAGAATTTAAATGAATCTCTCATCGATCTCCAAGAACTTGGAAAGTATGAGCAGTATATAAAATGGCCATGGTACATTTGGCTAGGTTTTATAGCTGGCTTGATTGCCATAGTAATGGTGACAATTATGCTTTGCTGTATGACCAGTTGCTGTAGTTGTCTCAAGGGCTGTTGTTCTTGTGGATCCTGCTGCAAATTTGATGAAGACGACTCTGAGCCAGTGCTCAAAGGAGTCAAATTACATTACACATAA

#### 3. Optimized codon sequence preferred by human hosts

ATGTTCGTCTTCCT CGTGCTGCTGCCTCTCGTCAGCAGTCAGTGTGTGAACCTCACTACTCGCACACAGCTGCCTCCAGCATACACTAATTCC TCACTAGAGGAGTCTACTATCCAGATAAGGTCTTCAGATCATCTGTGCTGCACAGCACACAGGATCTCTTCCTTCCATTCTTCAGCAATGTCACTTGGTTCCATGCTATACATGTGTCTGGCACCAATGGCACCAAGAGATTCGACAATCCTGTGCTGCCATTCAATGATGGTGTCTACTTCGCCAGCACAGAGAAGAGCAACATCATCAGAGGCTGGATCTTCGGAACTACACTAGATAGTAAGACACAGTCACTGCTCATCGTGAACAATGCCACCAATGTAGTGATCAAGGTGTGTGAGTTCCAGTTCTGCAATGATCCATTCCTTGGTGTCTACTATCATAAGAATAACAAGAGCTGGATGGAGAGTGAGTTCAGAGTCTACAGCAGTGCTAACAACTGTACATTCGAGTATGTATCTCAGCCATTCCTGATGGATCTGGAAGGCAAGCAAGGCAACTTCAAGAACCTGAGAGAGTTCGTGTTCAAGAATATAGATGGTTACTTCAAGATCTACAGCAAGCATACACCAATTAATCTAGTGAGAGATCTGCCTCAAGGCTTCAGTGCTCTAGAGCCACTAGTAGATCTGCCTATTGGCATCAACATCACTCGATTCCAGACACTACTAGCTCTGCACAGATCATATCTCACTCCTGGAGACAGCAGCAGTGGCTGGACAGCTGGTGCTGCTGCATACTATGTAGGATATCTGCAGCCAAGGACCTTCCTGCTGAAGTATAATGAGAATGGAACTATCACAGATGCTGTAGACTGTGCTCTAGATCCTCTCAGTGAGACTAAGTGTACTCTGAAGTCCTTCACTGTAGAGAAGGGCATCTATCAGACATCTAACTTCAGAGTGCAGCCAACAGAGAGCATCGTGCGCTTCCCTAACATCACTAACCTCTGTCCATTCGGAGAAGTCTTCAATGCTACTCGCTTCGCATCTGTCTATGCATGGAACAGGAAGCGCATCAGCAACTGTGTAGCTGACTACAGTGTGCTGTACAACAGTGCTAGCTTCAGCACATTCAAGTGCTATGGTGTGTCTCCAACCAAGCTCAATGATCTGTGCTTCACCAATGTCTATGCTGACAGCTTCGTCATCAGAGGAGATGAGGTGAGACAGATAGCTCCTGGACAGACAGGCAAGATAGCTGACTATAATTACAAGCTTCCTGATGACTTCACTGGCTGCGTCATAGCATGGAACAGCAACAATCTAGATAGTAAGGTTGGTGGCAACTATAATTATCTCTACAGACTCTTCAGGAAGAGCAACCTGAAGCCATTCGAGAGAGACATCAGCACAGAGATCTATCAGGCTGGAAGCACACCATGTAATGGTGTGGAAGGCTTCAACTGCTACTTCCCTCTGCAGAGCTATGGCTTCCAGCCAACTAATGGAGTTGGATATCAGCCATATCGAGTAGTAGTGCTGAGCTTCGAGCTGCTGCACGCTCCTGCTACAGTATGTGGACCTAAGAAGAGCACTAATCTAGTGAAGAACAAGTGTGTGAACTTCAACTTCAATGGACTCACTGGAACTGGAGTGCTGACAGAGAGCAACAAGAAGTTCCTGCCATTCCAGCAGTTCGGAAGAGATATAGCAGATACTACAGATGCAGTGAGAGATCCACAGACACTAGAGATACTAGACATCACTCCTTGCAGCTTCGGAGGAGTGTCTGTGATCACTCCTGGAACTAATACATCTAATCAAGTAGCTGTGCTGTATCAGGATGTCAACTGCACAGAGGTGCCAGTAGCTATACATGCAGATCAGCTGACTCCAACATGGAGAGTCTACAGCACAGGCAGCAATGTCTTCCAGACTAGAGCTGGCTGTCTCATAGGAGCAGAGCATGTCAATAATTCATATGAGTGTGACATACCAATTGGTGCTGGCATCTGTGCTAGCTATCAGACACAGACTAATTCTCCAAGACGAGCTAGATCAGTAGCATCTCAGAGCATCATAGCATATACTATGTCTCTAGGAGCTGAGAATTCAGTAGCATACAGCAACAACAGCATAGCTATACCAACCAACTTCACTATATCTGTGACTACAGAGATACTGCCTGTCAGCATGACTAAGACATCAGTAGACTGTACTATGTACATCTGTGGAGATAGTACAGAGTGCAGCAACCTGCTGCTGCAGTATGGAAGCTTCTGCACACAGCTGAACAGAGCTCTGACTGGAATAGCAGTAGAGCAGGACAAGAATACACAGGAGGTGTTCGCTCAAGTGAAGCAGATCTACAAGACACCTCCAATTAAGGACTTCGGTGGCTTCAACTTCTCTCAGATACTGCCAGATCCTTCCAAGCCTTCCAAGCGCAGCTTCATCGAGGATCTGCTCTTCAACAAGGTCACTCTAGCAGATGCTGGCTTCATCAAGCAGTATGGAGACTGTCTAGGAGACATAGCAGCTAGAGATCTCATCTGTGCTCAGAAGTTCAATGGCCTGACTGTGCTGCCTCCTCTGCTGACAGATGAGATGATAGCACAGTACACATCAGCTCTGCTAGCAGGAACTATCACTAGTGGCTGGACCTTCGGTGCTGGAGCTGCTCTGCAGATACCATTCGCGATGCAGATGGCATATAGATTCAATGGAATTGGAGTGACTCAGAATGTGCTCTATGAGAATCAGAAGCTGATAGCTAACCAGTTCAATTCAGCAATTGGCAAGATACAGGACTCTCTGAGCAGCACAGCATCTGCACTAGGCAAGCTGCAGGATGTCGTCAACCAGAATGCACAAGCACTCAACACTCTAGTGAAGCAGCTGAGCAGCAACTTCGGTGCTATATCATCTGTGCTGAATGACATACTGAGCAGACTAGACAAGGTAGAGGCAGAGGTGCAGATCGACAGACTCATCACTGGAAGACTGCAGTCTCTGCAGACATATGTCACTCAGCAGCTCATCAGAGCTGCTGAGATCAGAGCATCTGCTAATCTAGCAGCTACTAAGATGTCTGAGTGTGTGCTAGGCCAGAGCAAGAGAGTAGACTTCTGTGGCAAGGGCTATCATCTGATGAGCTTCCCTCAGAGTGCTCCTCATGGCGTCGTCTTCCTGCATGTGACATATGTGCCTGCTCAGGAGAAGAACTTCACTACAGCACCTGCTATCTGTCATGATGGCAAGGCTCACTTCCCAAGAGAAGGAGTCTTCGTGTCTAATGGCACACACTGGTTCGTGACTCAGCGCAACTTCTATGAGCCTCAGATCATCACTACAGATAACACCTTCGTGTCTGGCAACTGTGATGTCGTCATCGGCATCGTGAATAATACTGTCTATGATCCACTGCAGCCTGAGCTAGATAGCTTCAAGGAGGAGCTAGATAAGTACTTCAAGAACCACACATCTCCAGATGTAGATCTAGGAGACATCAGTGGAATTAATGCATCTGTCGTCAATATACAGAAGGAGATAGACAGACTCAATGAAGTGGCCAAGAACTTGAATGAGTCTCTGATAGATCTGCAGGAACTTGGCAAGTATGAGCAGTACATCAAGTGGCCTTGGTACATCTGGCTTGGATTCATAGCTGGCCTCATCGCTATCGTGATGGTGACTATCATGCTGTGCTGCATGACATCATGCTGCAGCTGTCTGAAGGGCTGCTGCAGCTGCGGCAGCTGCTGTAAGTTCGACGAGGATGACAGTGAGCCTGTGCTGAAGGGTGTGAAGCTGCACTACACA

### A.2 Insulin

#### 1 Protein sequence

NP 000198.1 INS [organism=Homo sapiens] [GeneID=3630]

MALWMRLLPLLALLALWGPDPAAAFVNQHLCGSHLVEALYLVCGERGFFYTPKT
RREAEDLQVGQVELGGGPGAGSLQPLALEGSLQKRGIVEQCCTSICSLYQLENYCN

#### 2. Nucleotide sequence

NC 000011.10:c2161209-2159779 INS [organism=Homo sapiens] [GeneID=3630] [chromosome=11]

ATGGCCCTGTGGATGCGCCTCCTGCCCCTGCTGGCGCTGCTGGCCCTCTGGGGACCTGACCCAGCCGCAGCCTTTGTGAACCAACACCTGTGCGGCTCACACCTGGTGGAAGCTCTCTACCTAGTGTGCGGGGAACGAGGCTTCTTCTACACACCCAAGACCCGCCGGGAGGCAGAGGACCTGCAGGTGGGGCAGGTGGAGCTGGGCGGGGGCCCTGGTGCAGGCAGCCTGCAGCCCTTGGCCCTGGAGGGGTCCCTGCAGAAGCGTGGCATTGTGGAACAATGCTGTACCAGCATCTGCTCCCTCTACCAGCTGGAGAACTACT

GCAACTAG

#### 3. Optimized codon sequence preferred by *E. coli* hosts

ATGGCGCTATGGATGCGACTACTACCACTACTCGCACTACTCGCACTATGGGGACCAGATCCTGCTGCTGCATTCGTTAATCAGCATCTCTGTGGTTCTCATCTCGTCGAGGCGCTATATCTCGTCTGTGGTGAGAGAGGATTCTTCTACACACCTAAGACACGACGAGAGGCAGAGGATCTACAGGTTGGTCAGGTAGAGCTCGGTGGTGGACCAGGAGCAGGATCGCTACAGCCACTAGCGCTGGAAGGTTCACTACAGAAGAGAGGAATAGTAGAGCAGTGCTGTACATCGATATGCTCACTATATCAGCTCGAGAATTACTGTAAT

## References

Agashe and Shankar, 2014.Agashe, D. and Shankar, N. (2014). The evolution of bacterial dna base composition. Journal of Experimental Zoology Part B: Molecular and Developmental Evolution, 322(7):517–528.

Albash and Lidar, 2018.Albash, T. and Lidar, D. A. (2018). Demonstration of a Scaling Advantage for a Quantum Annealer over Simulated Annealing. Physical Review X, 8(3):031016.

Alexaki et al., 2019.Alexaki, A., Kames, J., Holcomb, D. D., Athey, J., Santana-Quintero, L. V., Lam, P. V. N., Hamasaki-Katagiri, N., Osipova, E., Simonyan, V., Bar, H., Komar, A. A., and Kimchi-Sarfaty, C. (2019). Codon and codon-pair usage tables (cocoputs): Facilitating genetic variation analyses and recombinant gene design. Journal of Molecular Biology, 431(13):2434–2441. Computation Resources for Molecular Biology.

Andersson and Kurland, 1990.Andersson, S. G. and Kurland, C. G. (1990). Codon preferences in free-living microorganisms. Microbiol. Rev., 54(2):198–210.

Apolloni et al., 1990.Apolloni, B., Cesa-Bianchi, N., and De Falco, D. (1990). A numerical implementation of ”quantum annealing”. In Stochastic Processes, Physics and Geometry: Proceedings of the Ascona-Locarno Conference, pages 97–111.

Baeshen et al., 2014.Baeshen, N. A., Baeshen, M. N., Sheikh, A., Bora, R. S., Ahmed, M. M. M., Ramadan, H. A. I., Saini, K. S., and Redwan, E. M. (2014). Cell factories for insulin production. Microb. Cell Fact., 13(1):141.

Bennetzen and Hall, 1982.Bennetzen, J. L. and Hall, B. D. (1982). Codon selection in yeast. Journal of Biological Chemistry, 257(6):3026–3031.

Berkhout and van Hemert, 2015.Berkhout, B. and van Hemert, F. (2015). On the biased nucleotide composition of the human coronavirus RNA genome. Virus Res., 202:41–47.

Bharti et al., 2022.Bharti, K., Cervera-Lierta, A., Kyaw, T. H., Haug, T., Alperin-Lea, S., Anand, A., Degroote, M., Heimonen, H., Kottmann, J. S., Menke, T., Mok, W.-K., Sim, S., Kwek, L.-C., and Aspuru-Guzik, A. (2022). Noisy intermediate-scale quantum algorithms. Reviews of Modern Physics, 94(1):015004.

Buhr et al., 2016.Buhr, F., Jha, S., Thommen, M., Mittelstaet, J., Kutz, F., Schwalbe, H., Rodnina, M. V., and Komar, A. A. (2016). Synonymous Codons Direct Co-translational Folding toward Different Protein Conformations. Molecular Cell, 61(3):341–351.

Bulmer, 1987.Bulmer, M. (1987). Coevolution of codon usage and transfer RNA abundance. Nature, 325(6106):728–730.

Burgess-Brown et al., 2008.Burgess-Brown, N. A., Sharma, S., Sobott, F., Loenarz, C., Oppermann, U., and Gileadi, O. (2008). Codon optimization can improve expression of human genes in escherichia coli: A multi-gene study. Protein Expression and Purification, 59(1):94–102.

Cerezo et al., 2021.Cerezo, M., Arrasmith, A., Babbush, R., Benjamin, S. C., Endo, S., Fujii, K., McClean, J. R., Mitarai, K., Yuan, X., Cincio, L., and Coles, P. J. (2021). Variational quantum algorithms. Nature Reviews Physics, 3(9):625–644.

Clarke and Clark, 2008.Clarke, IV, T. F. and Clark, P. L. (2008). Rare codons cluster. PLOS ONE, 3(10):1–5.

Coghlan and Wolfe, 2000.Coghlan, A. and Wolfe, K. H. (2000). Relationship of codon bias to mRNA concentration and protein length in saccharomyces cerevisiae. Yeast, 16(12):1131–1145.

Coleman et al., 2008.Coleman, J. R., Papamichail, D., Skiena, S., Futcher, B., Wimmer, E., and Mueller, S. (2008). Virus Attenuation by Genome-Scale Changes in Codon Pair Bias. Science, 320(5884):1784–1787.

D-Wave Systems Inc., 2021.D-Wave Systems Inc. (2021). Hybrid Solver for Constrained Quadratic Models [White paper]. Technical report.

Deb and Jain, 2014.Deb, K. and Jain, H. (2014). An evolutionary many-objective optimization algorithm using reference-point-based nondominated sorting approach, part i: Solving problems with box constraints. IEEE Transactions on Evolutionary Computation, 18(4):577–601.

Djidjev, 2023.Djidjev, H. N. (2023). Logical qubit implementation for quantum annealing: augmented Lagrangian approach. Quantum Science and Technology, 8(3):035013.

Fox et al., 2021.Fox, D. M., Branson, K. M., and Walker, R. C. (2021). mRNA codon optimization with quantum computers. PLOS ONE, 16(10):e0259101.

Fu et al., 2013.Fu, Z., Gilbert, E. R., and Liu, D. (2013). Regulation of insulin synthesis and secretion and pancreatic beta-cell dysfunction in diabetes. Curr. Diabetes Rev., 9(1):25–53.

Fuglsang, 2006.Fuglsang, A. (2006). Estimating the ”Effective Number of Codons”: The Wright Way of Determining Codon Homozygosity Leads to Superior Estimates. Genetics, 172(2):1301–1307.

Gabbassov et al., 2023.Gabbassov, E., Rosenberg, G., and Scherer, A. (2023). Quantum Optimization: Lagrangian Dual versus QUBO in Solving Constrained Problems. arXiv.

Gerashchenko et al., 2021.Gerashchenko, M. V., Peterfi, Z., Yim, S. H., and Gladyshev, V. N. (2021). Translation elongation rate varies among organs and decreases with age. Nucleic Acids Res., 49(2):e9.

Grantham et al., 1980a.Grantham, R., Gautier, C., and Gouy, M. (1980a). Codon frequencies in 119 individual genes confirm consistent choices of degenerate bases according to genome type. Nucleic Acids Res., 8(9):1893–1912.

Grantham et al., 1980b.Grantham, R., Gautier, C., Gouy, M., Mercier, R., and Pavé, A. (1980b). Codon catalog usage and the genome hypothesis. Nucleic Acids Research, 8(1):197–197.

Gu et al., 2021.Gu, Q., Wang, R., Xie, H., Li, X., Jiang, S., and Xiong, N. (2021). Modified non-dominated sorting genetic algorithm iii with fine final level selection. Applied Intelligence, 51(7):4236–4269.

Gupta et al., 2017.Gupta, V., Sengupta, M., Prakash, J., and Tripathy, B. C. (2017). Production of recombinant pharmaceutical proteins. In Basic and Applied Aspects of Biotechnology, pages 77–101. Springer Singapore, Singapore.

Gustafsson et al., 2004.Gustafsson, C., Govindarajan, S., and Minshull, J. (2004). Codon bias and heterologous protein expression. Trends in Biotechnology, 22(7):346–353.

Hauke et al., 2020.Hauke, P., Katzgraber, H. G., Lechner, W., Nishimori, H., and Oliver, W. D. (2020). Perspectives of quantum annealing: methods and implementations. Reports on Progress in Physics, 83(5):054401.

Hia et al., 2019.Hia, F., Yang, S. F., Shichino, Y., Yoshinaga, M., Murakawa, Y., Vandenbon, A., Fukao, A., Fujiwara, T., Landthaler, M., Natsume, T., Adachi, S., Iwasaki, S., and Takeuchi, O. (2019). Codon bias confers stability to human mRNAs. EMBO Rep., 20(11):e48220.

Hiraoka et al., 2009.Hiraoka, Y., Kawamata, K., Haraguchi, T., and Chikashige, Y. (2009). Codon usage bias is correlated with gene expression levels in the fission yeast schizosaccharomyces pombe. Genes Cells, 14(4):499–509.

Holm, 1986.Holm, L. (1986). Codon usage and gene expression. Nucleic Acids Research, 14(7):3075–3087.

Ikemura, 1981.Ikemura, T. (1981). Correlation between the abundance of Escherichia coli transfer RNAs and the occurrence of the respective codons in its protein genes: A proposal for a synonymous codon choice that is optimal for the E. coli translational system. Journal of Molecular Biology, 151(3):389–409.

Ikemura, 1985.Ikemura, T. (1985). Codon usage and tRNA content in unicellular and multicellular organisms. Mol. Biol. Evol., 2(1):13–34.

Kadowaki and Nishimori, 1998.Kadowaki, T. and Nishimori, H. (1998). Quantum annealing in the transverse Ising model. Physical Review E, 58(5):5355–5363. First Suggestion about Quantum Annealing.

Kandeel et al., 2020.Kandeel, M., Ibrahim, A., Fayez, M., and Al-Nazawi, M. (2020). From SARS and MERS CoVs to SARS-CoV-2: Moving toward more biased codon usage in viral structural and nonstructural genes. J. Med. Virol., 92(6):660–666.

Kane, 1995.Kane, J. F. (1995). Effects of rare codon clusters on high-level expression of heterologous proteins in escherichia coli. Current Opinion in Biotechnology, 6(5):494–500.

Karimi and Ronagh, 2017.Karimi, S. and Ronagh, P. (2017). A subgradient approach for constrained binary optimization via quantum adiabatic evolution. Quantum Information Processing, 16(8):185.

Karp, 1972.Karp, R. M. (1972). Reducibility among Combinatorial Problems, pages 85–103. Springer US, Boston, MA.

Khattak et al., 2021.Khattak, S., Rauf, M. A., Zaman, Q., Ali, Y., Fatima, S., Muhammad, P., Li, T., Khan, H. A., Khan, A. A., Ngowi, E. E., Wu, D.-D., and Ji, X.-Y. (2021). Genome-wide analysis of codon usage patterns of SARS-CoV-2 virus reveals global heterogeneity of COVID-19. Biomolecules, 11(6):912.

King et al., 2021.King, A. D., Raymond, J., Lanting, T., Isakov, S. V., Mohseni, M., Poulin-Lamarre, G., Ejtemaee, S., Bernoudy, W., Ozfidan, I., Smirnov, A. Y., Reis, M., Altomare, F., Babcock, M., Baron, C., Berkley, A. J., Boothby, K., Bunyk, P. I., Christiani, H., Enderud, C., Evert, B., Harris, R., Hoskinson, E., Huang, S., Jooya, K., Khodabandelou, A., Ladizinsky, N., Li, R., Lott, P. A., MacDonald, A. J. R., Marsden, D., Marsden, G., Medina, T., Molavi, R., Neufeld, R., Norouzpour, M., Oh, T., Pavlov, I., Perminov, I., Prescott, T., Rich, C., Sato, Y., Sheldan, B., Sterling, G., Swenson, L. J., Tsai, N., Volkmann, M. H., Whittaker, J. D., Wilkinson, W., Yao, J., Neven, H., Hilton, J. P., Ladizinsky, E., Johnson, M. W., and Amin, M. H. (2021). Scaling advantage over path-integral Monte Carlo in quantum simulation of geometrically frustrated magnets. Nature Communications, 12(1):1113.

Kirkpatrick et al., 1983.Kirkpatrick, S., Gelatt, Jr, C. D., and Vecchi, M. P. (1983). Optimization by simulated annealing. Science, 220(4598):671–680.

Koblan et al., 2018.Koblan, L. W., Doman, J. L., Wilson, C., Levy, J. M., Tay, T., Newby, G. A., Maianti, J. P., Raguram, A., and Liu, D. R. (2018). Improving cytidine and adenine base editors by expression optimization and ancestral reconstruction. Nat. Biotechnol., 36(9):843–846.

Kudla et al., 2006.Kudla, G., Lipinski, L., Caffin, F., Helwak, A., and Zylicz, M. (2006). High guanine and cytosine content increases mRNA levels in mammalian cells. PLoS Biol., 4(6):e180.

Lehmann and Libchaber, 2008.Lehmann, J. and Libchaber, A. (2008). Degeneracy of the genetic code and stability of the base pair at the second position of the anticodon. RNA, 14(7):1264–1269.

Letzring et al., 2010.Letzring, D. P., Dean, K. M., and Grayhack, E. J. (2010). Control of translation efficiency in yeast by codon-anticodon interactions. RNA, 16(12):2516–2528.

Lewis and Brubaker, 2021.Lewis, G. F. and Brubaker, P. L. (2021). The discovery of insulin revisited: lessons for the modern era. J. Clin. Invest., 131(1).

Li et al., 2020.Li, X., Geng, M., Peng, Y., Meng, L., and Lu, S. (2020). Molecular immune pathogenesis and diagnosis of covid-19. Journal of Pharmaceutical Analysis, 10(2):102–108.

Liachko et al., 2014.Liachko, I., Youngblood, R. A., Tsui, K., Bubb, K. L., Queitsch, C., Raghuraman, M. K., Nislow, C., Brewer, B. J., and Dunham, M. J. (2014). GC-rich DNA elements enable replication origin activity in the methylotrophic yeast pichia pastoris. PLoS Genet., 10(3):e1004169.

Liao et al., 2023.Liao, X., Zhu, W., Zhou, J., Li, H., Xu, X., Zhang, B., and Gao, X. (2023). Repetitive DNA sequence detection and its role in the human genome. Commun. Biol., 6(1):954.

Liu, 2020.Liu, Y. (2020). A code within the genetic code: codon usage regulates co-translational protein folding. Cell Commun. Signal., 18(1):145.

Liu et al., 2023.Liu, Y., Liang, N., Xian, Q., and Zhang, W. (2023). GC heterogeneity reveals sequence-structures evolution of angiosperm ITS2. BMC Plant Biol., 23(1):608.

Lloyd and Sharp, 1991.Lloyd, A. T. and Sharp, P. M. (1991). Codon usage in aspergillus nidulans. Mol. Gen. Genet., 230(1-2):288–294.

Lucas, 2014.Lucas, A. (2014). Ising formulations of many NP problems. Frontiers in Physics, 2:5.

Lynch and Marinov, 2017.Lynch, M. and Marinov, G. K. (2017). Membranes, energetics, and evolution across the prokaryote-eukaryote divide. Elife, 6.

Menzella, 2011.Menzella, H. G. (2011). Comparison of two codon optimization strategies to enhance recombinant protein production in Escherichia coli. Microbial Cell Factories, 10(1):15.

Mitra et al., 2016.Mitra, S., Ray, S. K., and Banerjee, R. (2016). Synonymous codons influencing gene expression in organisms. Res. Rep. Biochem., 6:57–65.

Nakamura et al., 2000.Nakamura, Y., Gojobori, T., and Ikemura, T. (2000). Codon usage tabulated from international dna sequence databases: status for the year 2000. Nucleic acids research, 28(1):292–292. Database available from: https://www.kazusa.or.jp/codon/.

Obradovic et al., 2013.Obradovic, J., Jurisic, V., Tosic, N., Mrdjanovic, J., Perin, B., Pavlovic, S., and Djordjevic, N. (2013). Optimization of PCR conditions for amplification of GC-Rich EGFR promoter sequence. J. Clin. Lab. Anal., 27(6):487–493.

Ohzeki, 2020.Ohzeki, M. (2020). Breaking limitation of quantum annealer in solving optimization problems under constraints. Scientific Reports, 10(1):3126.

Palidwor et al., 2010.Palidwor, G. A., Perkins, T. J., and Xia, X. (2010). A general model of codon bias due to gc mutational bias. PLOS ONE, 5(10):1–11.

Parvathy et al., 2022.Parvathy, S. T., Udayasuriyan, V., and Bhadana, V. (2022). Codon usage bias. Mol. Biol. Rep., 49(1):539–565.

Phaniendra et al., 2015.Phaniendra, A., Jestadi, D. B., and Periyasamy, L. (2015). Free radicals: properties, sources, targets, and their implication in various diseases. Indian J. Clin. Biochem., 30(1):11–26.

Pozzoli et al., 2008.Pozzoli, U., Menozzi, G., Fumagalli, M., Cereda, M., Comi, G. P., Cagliani, R., Bresolin, N., and Sironi, M. (2008). Both selective and neutral processes drive GC content evolution in the human genome. BMC Evol. Biol., 8(1):99.

Preskill, 2018.Preskill, J. (2018). Quantum Computing in the NISQ era and beyond. Quantum, 2:79.

Puigbò et al., 2007.Puigbò, P., Guzmán, E., Romeu, A., and Garcia-Vallvé, S. (2007). OPTIMIZER: a web server for optimizing the codon usage of DNA sequences. Nucleic Acids Res., 35(Web Server issue):W126–31.

Rao et al., 2013.Rao, Y. S., Chai, X. W., Wang, Z. F., Nie, Q. H., and Zhang, X. Q. (2013). Impact of GC content on gene expression pattern in chicken. Genet. Sel. Evol., 45(1):9.

Redwan et al., 2008.Redwan, E.-R. M., Matar, S. M., El-Aziz, G. A., and Serour, E. A. (2008). Synthesis of the human insulin gene: protein expression, scaling up and bioactivity. Prep. Biochem. Biotechnol., 38(1):24–39.

Reeves and Rowe, 2002.Reeves, C. R. and Rowe, J. E. (2002). Genetic algorithms: Principles and perspectives. Operations Research/Computer Science Interfaces Series. Springer, New York, NY, 2002 edition.

Ronagh et al., 2016.Ronagh, P., Woods, B., and Iranmanesh, E. (2016). Solving constrained quadratic binary problems via quantum adiabatic evolution. Quantum Information & Computation, 16(11-12):1029–1047.

Rosano and Ceccarelli, 2009.Rosano, G. L. and Ceccarelli, E. A. (2009). Rare codon content affects the solubility of recombinant proteins in a codon bias-adjusted escherichia coli strain. Microb. Cell Fact., 8(1):41.

Rosano and Ceccarelli, 2014.Rosano, G. L. and Ceccarelli, E. A. (2014). Recombinant protein expression in escherichia coli: advances and challenges. Frontiers in Microbiology, 5.

S. K. Gupta and Ghosh, 2004.S. K. Gupta, T. K. B. and Ghosh, T. C. (2004). Synonymous codon usage in lactococcus lactis: Mutational bias versus translational selection. Journal of Biomolecular Structure and Dynamics, 21(4):527–535. PMID: 14692797.

Sharp et al., 1988.Sharp, P. M., Cowe, E., Higgins, D. G., Shields, D. C., Wolfe, K. H., and Wright, F. (1988). Codon usage patterns in escherichia coli, bacillus subtilis, saccharomyces cerevisiae, schizosaccharomyces pombe, drosophila melanogaster and homo sapiens; a review of the considerable within-species diversity. Nucleic Acids Res., 16(17):8207–8211.

Sharp and Li, 1986.Sharp, P. M. and Li, W. H. (1986). An evolutionary perspective on synonymous codon usage in unicellular organisms. J. Mol. Evol., 24(1-2):28–38.

Sharp and Li, 1987.Sharp, P. M. and Li, W.-H. (1987). The codon adaptation index-a measure of directional synonymous codon usage bias, and its potential applications. Nucleic Acids Research, 15(3):1281–1295.

Sharp and Lloyd, 1993.Sharp, P. M. and Lloyd, A. T. (1993). Regional base composition variation along yeast chromosome III: evoluation of chormosome primary structure. Nucleic Acids Research, 21(2):179–183.

Shi et al., 2022.Shi, A., Fan, F., and Broach, J. R. (2022). Microbial adaptive evolution. J. Ind. Microbiol. Biotechnol., 49(2).

Shields and Sharp, 1987.Shields, D. C. and Sharp, P. M. (1987). Synonymous codon usage in bacillus subtilis reflects both translational selection and mutational biases. Nucleic Acids Res., 15(19):8023–8040.

Simmonds, 2012.Simmonds, P. (2012). SSE: a nucleotide and amino acid sequence analysis platform. BMC Res. Notes, 5(1):50.

Suspène et al., 2008.Suspène, R., Renard, M., Henry, M., Guétard, D., Puyraimond-Zemmour, D., Billecocq, A., Bouloy, M., Tangy, F., Vartanian, J.-P., and Wain-Hobson, S. (2008). Inversing the natural hydrogen bonding rule to selectively amplify GC-rich ADAR-edited RNAs. Nucleic Acids Res., 36(12):e72.

Trigiante et al., 2021.Trigiante, G., Blanes Ruiz, N., and Cerase, A. (2021). Emerging roles of repetitive and repeat-containing rna in nuclear and chromatin organization and gene expression. Frontiers in Cell and Developmental Biology, 9.

Victor et al., 2019.Victor, M. P., Acharya, D., Begum, T., and Ghosh, T. C. (2019). The optimization of mRNA expression level by its intrinsic properties-insights from codon usage pattern and structural stability of mRNA. Genomics, 111(6):1292–1297.

Wheeler et al., 2007.Wheeler, D. L., Barrett, T., Benson, D. A., Bryant, S. H., Canese, K., Chetvernin, V., Church, D. M., DiCuccio, M., Edgar, R., Federhen, S., et al. (2007). Database resources of the national center for biotechnology information. Nucleic acids research, 36(suppl 1):D13–D21. Database available from: https://www.ncbi.nlm.nih.gov (accessed on 16 February 2021).

Wright, 1990.Wright, F. (1990). The ‘effective number of codons’ used in a gene. Gene, 87(1):23–29.

Xu et al., 2008.Xu, X.-Z., Liu, Q.-P., Fan, L.-J., Cui, X.-F., and Zhou, X.-P. (2008). Analysis of synonymous codon usage and evolution of begomoviruses. J. Zhejiang Univ. Sci. B, 9(9):667–674.

Yarkoni et al., 2022.Yarkoni, S., Raponi, E., Bäck, T., and Schmitt, S. (2022). Quantum annealing for industry applications: introduction and review. Reports on Progress in Physics, 85(10):104001.

Yokobayashi et al., 2002.Yokobayashi, Y., Weiss, R., and Arnold, F. H. (2002). Directed evolution of a genetic circuit. Proceedings of the National Academy of Sciences, 99(26):16587–16591.

Yoshikawa and Sueoka, 1963.Yoshikawa, H. and Sueoka, N. (1963). Sequential replication of bacillus subtilis chromosome, i. comparison of marker frequencies in exponential and stationary growth phases. Proceedings of the National Academy of Sciences, 49(4):559–566.

Zhang et al., 2024.Zhang, H., Sarkar, A., and Bertels, K. (2024). A resource-efficient variational quantum algorithm for mRNA codon optimization. arXiv.

Šmarda et al., 2014.Šmarda, P., Bureš, P., Horová, L., Leitch, I. J., Mucina, L., Pacini, E., Tichý, L., Grulich, V., and Rotreklová, O. (2014). Ecological and evolutionary significance of genomic gc content diversity in monocots. Proceedings of the National Academy of Sciences, 111(39):E4096–E4102.

